# Canonical and phosphoribosyl ubiquitination coordinate to stabilize a proteinaceous structure surrounding the *Legionella*-containing vacuole

**DOI:** 10.1101/2025.07.22.666189

**Authors:** Adriana Steinbach, Chetan Mokkapati, Puspangana Singh, Shaeri Mukherjee

## Abstract

*Legionella pneumophila* (*L.p.*), an intracellular bacterial pathogen, hijacks the ubiquitin signaling network of its eukaryotic host cells to establish infection. Two families of *L.p.* secreted ubiquitin ligases are instrumental in the maturation of the *Legionella*-containing vacuole (LCV): the SidC/SdcA family, which catalyzes canonical ubiquitination, and the SidE family, which bypasses the E1-E2-E3 enzymatic cascade and directly conjugates ubiquitin to a target through a phosphoribosyl (PR) linkage. Here, we demonstrate that the coordinated activities of these two effector families generate a hyperstable, ubiquitin-rich structure surrounding the LCV. We propose a model in which an initial wave of SidC/SdcA-mediated canonical ubiquitination around the LCV is further modified by SidE family-driven PR-ubiquitination, resulting in a detergent-resistant “cloud”. The “cloud” is transient, breaking down as infection progresses, suggesting that *L.p.* reshapes the properties of the proteinaceous shell surrounding the vacuole to meet changing needs throughout its intracellular lifecycle. This unusual structure likely stabilizes and protects the LCV, shielding it from host defense mechanisms during early infection. Our findings reveal cellular consequences of effector interplay during infection and provide a foundation for future studies into the structure and function of the proteinaceous “cloud” surrounding the LCV.

## INTRODUCTION

Protein ubiquitination is one of the most versatile signaling mechanisms in eukaryotic cells. Ubiquitin, a small, highly conserved globular protein, can be covalently linked to a target protein, serving as a post-translational modification. Most frequently, this linkage is an isopeptide bond between the carboxy terminus of ubiquitin and a lysine side chain or amino terminal amine group on the target protein. Ubiquitin itself can be ubiquitinated, resulting in polyubiquitin chains, and the identity of the residues used to form these intermolecular linkages dictates downstream signaling outcomes^1^. The modularity of the ubiquitin conjugation system, in which attachment site, chain length, and chain linkage pattern affect signaling and target protein activity, confers the versatility required to regulate a broad range of cellular processes. Examples include cell division and differentiation, protein homeostasis, autophagy, immune signaling, and membrane trafficking^2,3^. Given the critical role and flexibility of the “ubiquitin code”, it is unsurprising that many pathogens have evolved to subvert this signaling system as part of their infection strategy^4^.

*Legionella pneumophila* (*L.p.*) is an intracellular bacterial pathogen of diverse eukaryotic cells. In humans, *L.p.* infection of alveolar macrophages can cause a severe pneumonia known as Legionnaires’ disease; however, humans are a dead-end host for the bacterium. The natural hosts for *L.p.* are free-living protozoa across multiple phyla, demonstrating the bacterium’s remarkable ability to establish a protected niche within evolutionarily divergent eukaryotic cells^5^. Upon phagocytic uptake, *L.p.* uses a type IV secretion system (T4SS) to translocate over 300 bacterial proteins, known as effectors, across the phagosome membrane and into the host cell. These effectors regulate host processes on a global scale, such as protein translation^6,7^ and gene expression^8^, while simultaneously remodeling the phagosomal membrane surrounding the bacterium to create a specialized compartment known as the *Legionella*-containing vacuole (LCV)^9,10^. Isogenic strains lacking a functional T4SS are efficiently trafficked to the lysosome after internalization, underscoring the essential role of bacterial effectors in *L.p.* pathogenesis^11,12^.

Mounting evidence suggests that hijacking the host’s ubiquitin signaling network is an important element of *L.p.* pathogenesis^13^. Ubiquitinated proteins begin to accumulate around the LCV during early infection^14,15^, and over 20 bacterial effectors have been shown to act on the ubiquitin system in various ways^16^. In eukaryotic cells, ubiquitination is a highly regulated process involving a three-enzyme cascade: an E1 activating enzyme, an E2 conjugating enzyme, and an E3 ligase, which, in coordination with the E2, facilitates the covalent attachment of ubiquitin to a target protein. This process is reversible, as cells express deubiquitinating enzymes (DUBs) that hydrolyze the isopeptide bonds linking ubiquitin to substrates^1^. *L.p.* secretes multiple effectors that act as E3 enzymes and DUBs^16^. Furthermore, recent studies have shown that four homologous *L.p.* effectors known as the SidE family can bypass the canonical eukaryotic E1-E2-E3 ubiquitination cascade. These effectors instead catalyze the formation of a phosphoribosyl (PR) linkage between an arginine residue on ubiquitin and a serine residue on the target protein, a process termed PR-ubiquitination^17,18^. The activity of the SidE family of effectors has been implicated in the manipulation of host endoplasmic reticulum (ER) proteins^19,20^, alterations in host metabolism^21^, and evasion of autophagy^22–25^ during infection.

Observations from multiple studies suggest that the SidE family and the SidC/SdcA family, effectors that catalyze canonical ubiquitination^26^, regulate closely related processes during infection. Both effector families are essential for recruitment of ER membrane to the LCV shortly after formation, a hallmark of LCV maturation^19,26,27^. Furthermore, both families ubiquitinate multiple small GTPases in the Rab family, as demonstrated by our lab^28,29^ and others^17,30,31^. A recent study provides evidence that these small GTPases are modified with mixed canonical and PR-linked ubiquitin chains during infection through the combined activities of the SidC and SidE effector families, and that the SidE family of effectors may crosslink these chains^24^. This hypothesis aligns with the intriguing proposal that ER membrane surrounding the LCV during infection is stabilized by SidE family activity, forming a protective barrier around the vacuole^32^.

In this work, we seek to place the biochemical activities of the SidC and SidE effector families within the context of cell biological events at the LCV membrane. Using Rab5 as a model substrate—previously shown by our lab to require both the SidC and SidE effectors for maximal ubiquitination and LCV recruitment^28^—we demonstrate that the morphology of the Rab5-positive regions surrounding the LCV depends on SidE activity. Further, we observe that Rab5 forms a detergent-resistant structure around wild-type (WT) LCVs during early infection. We identify a similar pattern for ubiquitin itself, where SidE activity is required to establish the distinctive, cloud-like morphology and detergent resistance of ubiquitin associated with the LCV. We find that this hyperstable structure is disassembled as infection progresses, indicating that the ubiquitin shell may provide protection shortly after internalization. We propose a model in which the SidC family initiates a wave of canonical ubiquitination around the LCV early in infection, forming a tight, detergent-sensitive ubiquitin coat. This coat is subsequently modified by the SidE family, likely in tandem with further ubiquitin chain synthesis by SidC/SdcA^24^, creating an expansive, detergent-resistant structure. Live imaging of ubiquitin during infection supports this hypothesis, as ubiquitin surrounding WT LCVs expands rapidly outward from the bacterial cell, while in the SidE knockout strain, ubiquitin remains tightly LCV-associated. Further, timelapse imaging of both Rab5 and ubiquitin during infection demonstrates that LCV recruitment of these two host proteins is temporally coupled by the activity of the SidE family of effectors. Our findings highlight the complex interplay between the SidE and SidC effector families and provide a foundation for future studies on the physical and cell biological properties of the enigmatic proteinaceous structure surrounding the LCV.

## RESULTS

### Rab5 displays a distinctive “cloud” morphology when associated with the WT LCV

While it has previously been reported that the host small GTPase Rab5 is recruited to LCVs harboring the T4SS-null *dotA* strain^33^, our lab recently demonstrated that Rab5 also associates with the WT LCV throughout early infection. We found that maximal Rab5 association with WT LCVs requires the SidC/SdcA and SidE bacterial effector families, indicating that this process is driven by bacterial effector activity. In contrast, Rab5 recruitment to the *dotA* LCV likely occurs passively as part of endosome maturation^28^. We realized that we had serendipitously discovered in Rab5 a tool to study how proteins at the LCV are manipulated by bacterial effectors. While mitochondria make close contacts with both the WT and *dotA* LCV^34^, to our knowledge no host protein aside from Rab5 has been observed to localize to both the virulent and avirulent LCV at the same time point during infection. We began by examining immunofluorescence micrographs of HeLa FcγR infected with *L.p.* WT or *dotA* fixed at 1-hour post-infection (hpi) and stained with anti-Rab5 antibody. We observed that, while Rab5 signal closely conformed to the bacterial cell outline for *dotA* LCVs, WT LCVs exhibited a cloud-like Rab5 morphology that extended beyond the bacterial cell (**Fig. 1A**). To quantify LCV-associated Rab5 area, we selected Rab5-positive LCVs for both strains and used CellProfiler^35^ to measure the area of the bacterial cell and the LCV-associated Rab5-positive region (see Methods). Calculating the ratio of the Rab5-positive area to *L.p.* area to normalize for bacterial cell size, we found that, indeed, the Rab5-positive region surrounding the LCV was consistently larger for WT bacteria compared to *dotA* (**Fig. 1B**).

**Figure 1:**
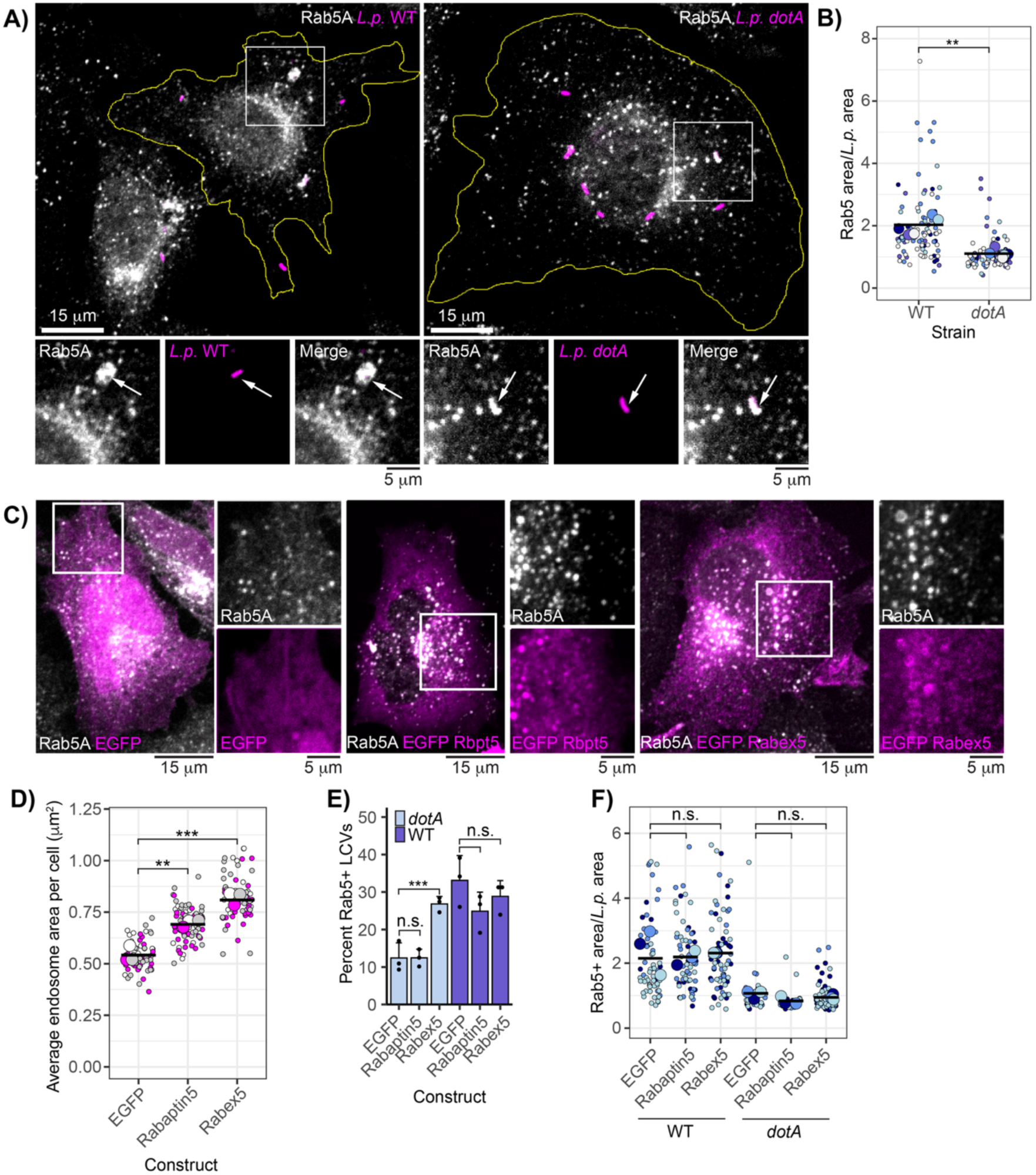
Rab5A forms a “cloud” around the WT LCV in a bacterial effector dependent manner. (A) Endogenous Rab5A recruitment to the WT or *dotA* LCV in HeLa FcγR. (B) LCV-associated Rab5A area normalized to bacterial cell area for WT and *dotA* strains. N=5, 8-30 LCVs measured per strain per replicate. Welch’s two-sample t-test, p = 0.0009. In this and all subsequent jitter plots, small points represent individual measurements, large points are mean values across a given biological replicate, and horizontal bars depict the sample mean across replicates. Color corresponds to biological replicate. (C) Endogenous Rab5A staining in cells overexpressing EGFP or indicated fusion constructs. (D) Mean endosome area per cell in samples overexpressing indicated construct. N=3, 20-40 cells scored per replicate. ANOVA, Tukey Kramer post hoc test, ** p<0.005, *** p<0.0005. (E) Manual scoring of endogenous Rab5 recruitment to WT or *dotA* LCVs in cells overexpressing indicated construct. N=3, 50-130 LCVs scored per replicate. G test, Bonferroni adj. p value = 0.0125, ***p<0.000125. (F) LCV-associated Rab5A area as in panel B in cells expressing indicated construct. N = 3, 10-30 LCVs scored per replicate. ANOVA, Tukey Kramer post hoc test, n.s. p>0.05. For all infection experiments, cells were fixed at 1hpi.

While we suspected that formation of the Rab5 cloud around the WT LCV is driven by bacterial effectors, it is possible that host regulators of Rab5 activity could contribute to this unusual morphology. Models include effector-induced hyperactivation of host regulators, or an effector-activated anti-pathogen defense mechanism mounted by the host. In mammalian cells, the switch between the inactive GDP-bound state and the active GTP-bound state for Rab5 is controlled by Rabex5, a guanine nucleotide exchange factor (GEF) specific to the Rab5 subfamily. Rabex5 forms a complex with Rabaptin5, which stimulates Rabex5 GEF activity and binds active Rab5, thereby creating a positive feedback loop that amplifies Rab5 activation at a cognate membrane ^36,37^. To assess whether either Rabex5 or Rabaptin5 promote Rab5 cloud formation, we conducted a series of overexpression experiments. In uninfected cells, overexpression of EGFP-Rabex5 or EGFP-Rabaptin5, compared to EGFP alone, resulted in Rab5-positive endosome swelling, a hallmark of Rab5 hyperactivation^38–40^, in agreement with previous results^41^ (**Fig. 1C-D**). Next, we infected cells overexpressing EGFP, EGFP-Rabaptin5, or EGFP-Rabex5 with either *L.p.* WT or *dotA* and assessed both LCV recruitment and morphology of endogenous Rab5. Rabex5 overexpression significantly increased Rab5 recruitment to the *dotA* LCV. Despite induction of endosome swelling, Rabaptin5 overexpression did not significantly increase Rab5 recruitment to the *dotA* LCV, perhaps suggesting that the cellular concentration of Rabex5 or some other factor is limiting in this context. Notably, neither construct altered Rab5 recruitment to WT LCVs (**Fig. 1E**). Furthermore, neither Rabex5 nor Rabaptin5 overexpression increased the LCV-associated Rab5 area for either strain (**Fig. 1F**). These results suggest that while excess Rabex5 hyperactivates Rab5, its overexpression does not significantly impact Rab5 recruitment or morphology at WT LCVs.

As these results suggest that Rab5 recruitment to the WT LCV may be driven by factors outside of the host cell’s regulation of Rab5’s GTPase cycle, we next assessed LCV localization of two Rab5 nucleotide-binding mutants during infection. Rab5 Q79L has reduced GTPase activity and is largely GTP-bound *in vivo*, whereas Rab5 S34N preferentially binds GDP, resulting in small early endosomes and a dominant negative effect on early endosome homotypic fusion ^38^. In cells transiently transfected with mCherry-fusion Rab5A variants, we observe LCV localization of S34N and Q79L at comparable levels during early infection (**Fig. S1AB**). Previously, we demonstrated that Rab5 is ubiquitinated during *L.p.* WT infection in a manner dependent on LCV localization^28^. Consistent with this finding, 3XFlag Rab5 S34N and Q79L are both ubiquitinated during *L.p.* WT infection, whereas a C-terminal truncation mutant (Rab5 1-211) lacking the prenylated CAAX motif required for membrane localization is not (**Fig. S1C**). Importantly, our lab has previously demonstrated that the CAAX truncation mutant is not recruited to the LCV during infection^28^. Taken together, these results suggest that Rab5 ubiquitination and recruitment to the LCV is not dependent upon nucleotide binding state, and that host regulators of Rab5 GTPase activity are not primary drivers of the unusual “cloud” morphology observed at the WT LCV, pointing instead to direct action on Rab5 by bacterial effectors.

### The SidE family of effectors is required for Rab5 cloud formation

Previously, our lab observed defects in Rab5 polyubiquitination and LCV recruitment during infection with an *L.p.* strain lacking the SidE family of effectors (*L.p.* Δ*sidE/sdeABC*) compared to *L.p.* WT^28^. Based on these results, we hypothesized that the SidE family might play a role in Rab5 cloud formation. While the majority of Δ*sidE/sdeABC* LCVs were Rab5-negative, 7+/- 3% (mean +/- SD) stained positive for Rab5 at 1hpi^28^. Closer examination of these Rab5-positive Δ*sidE/sdeABC* LCVs revealed tight Rab5 localization reminiscent of the *dotA* Rab5-positive LCVs (**Fig. 2A**). Next, we wanted to assess whether Rab5 cloud formation depends on the PR-ubiquitination activity of the SidE family. To this end, we generated an *L.p*. expression construct encoding SdeB with a non-functional mono-ADP ribosyltransferase (mART) domain by substituting the catalytic glutamate residues in the R-S-ExE motif for alanine (SdeB EE/AA)^17^. To verify loss of function for this mutant, we infected cells expressing ubiquitin lacking the final two C-terminal glycine residues (ubiquitin ΔGG), a substrate that canonical E3 ligases cannot utilize, but that can be incorporated into PR-linked polyubiquitin chains^20,24^. Infection with the SidE family knockout and the SdeB EE/AA complemented strain failed to induce ubiquitin ΔGG conjugation in cells, unlike infection with *L.p.* WT and the SdeB WT complemented knockout strain (**Fig. S2**). Finally, measuring Rab5 cloud formation around the LCV of the SidE family knockout strain panel, we found that plasmid-encoded SdeB WT, but not SdeB EE/AA, rescued Rab5 cloud formation (**Fig. 2AB**). These results suggest that SidE family-catalyzed PR-ubiquitination is necessary for Rab5 cloud formation.

**Figure 2:**
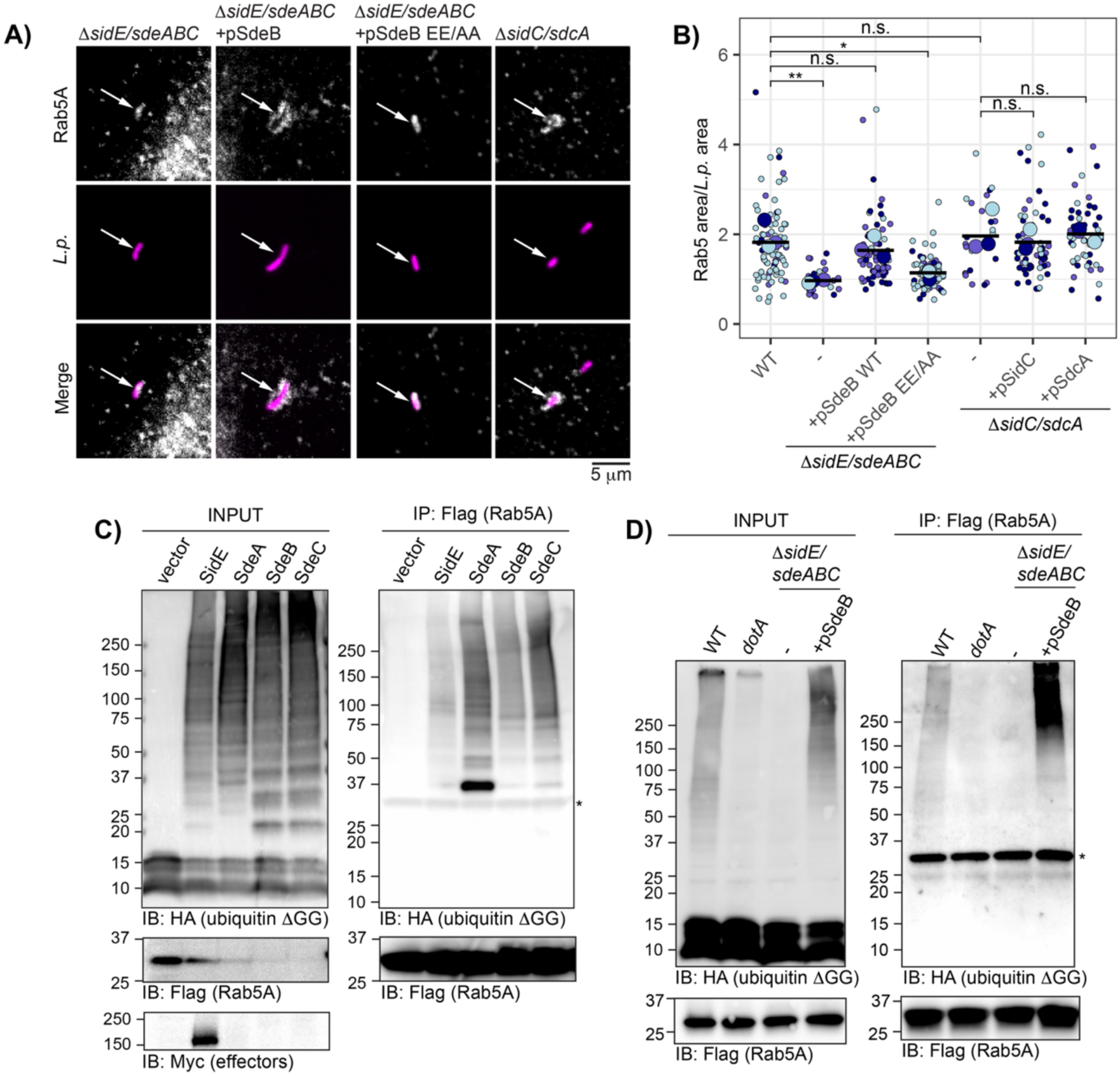
The SidE family is required for Rab5 cloud formation at the LCV. (A) Endogenous Rab5A recruitment to the LCV for the indicated strains in HeLa FcγR. (B) LCV-associated Rab5 area normalized to bacterial cell size for the indicated strains. N=3, 5-60 LCVs scored per strain per replicate. One way ANOVA followed by Tukey-Kramer post hoc test, n.s. p>0.05, * p<0.05, **p<0.005. (C) Immunoprecipitation of Flag-Rab5A from cells expressing HA ubiquitin 11GG and indicated myc-tagged SidE family effector, or vector control. (D) Immunoprecipitation of Flag-Rab5A from cells expressing HA ubiquitin 11GG and infected with indicated *L.p.* strain. For C and D, * indicates non-specific band, most likely light chain. For all infection experiments, cells were fixed or lysed at 1hpi.

Next, as we have reported that the SidC/SdcA canonical ligase effectors are also required for maximal Rab5 recruitment and ubiquitination during infection^28^, we assessed Rab5 cloud formation for the 7 +/- 2% Δ*sidC/sdcA* Rab5-positive LCVs observed at 1hpi^28^. Despite the strong defect in Rab5 recruitment for this strain, positive Δ*sidC/sdcA* LCVs showed clear Rab5 cloud formation, with normalized area measurements comparable to *L.p.* WT (**Fig. 2AB**). This finding suggests that SidC/SdcA may play a primary role in initial Rab5 recruitment or retention at the LCV, allowing further modification by the SidE family and subsequent cloud formation. The low frequency association of Rab5 with the Δ*sidC/sdcA* LCV may be driven by host or bacterial factors, or a combination of the two.

To verify that Rab5 is a substrate for SidE family-catalyzed PR-ubiquitination, we again utilized the ubiquitin ΔGG construct. First, to determine whether ectopic expression of the SidE family is sufficient for Rab5 PR-ubiquitination, we co-transfected cells with HA-ubiquitin 11GG, Flag-Rab5, and myc-tagged SidE family effectors (or vector alone). Immunoprecipitated Rab5 from cells expressing SidE family members exhibited HA signal across a range of molecular weights from approximately 37 kDa (monoubiquitinated Rab5A) to the top of the gel, consistent with extensive PR-ubiquitination. No HA-ubiquitin ΔGG conjugation was observed for the vector control (**Fig. 2C**). We note that expression of all effectors aside from SidE was extremely low, such that myc detection by Western blot was unsuccessful at the appropriate input loading volume, despite efficient HA-ubiquitin ΔGG conjugation in all effector transfected conditions. Finally, to assess PR-ubiquitination of Rab5 during infection, we co-transfected cells with HA-ubiquitin 11GG and Flag-Rab5, and infected with *L.p.* WT, *dotA*, or Δ*sidE/sdeABC* with and without SdeB complementation. High molecular weight HA signal was observed for Rab5 immunoprecipitated from cells infected with *L.p.* WT, but not *dotA* or Δ*sidE/sdeABC*. Complementation of the SidE family strain with plasmid-encoded SdeB WT robustly rescued the loss of HA-ubiquitin ΔGG conjugation to Rab5 (**Fig 2D**). Overall, our results indicate that PR-ubiquitination of Rab5 catalyzed by the SidE family of effectors is required for Rab5 cloud formation around the LCV, whereas SidC/SdcA likely contribute to initial recruitment of Rab5 to LCV during infection, where it can be further modified by the SidE family.

### Rab5 resists detergent washout when associated with the WT LCV

Having established that the Rab5-positive region around the WT LCV adopts a distinctive, cloud-like morphology in a SidE family-dependent manner, we sought to explore the properties of this structure. In our previous study, we found that polyubiquitinated Rab5 is enriched in the membrane fraction upon crude fractionation of *L.p.* infected cells, suggesting that polyubiquitination may increase stable association of Rab5 with membranes^28^. Additionally, it has been shown that upon SidE family mediated recruitment to the WT LCV, reticulon4 (Rtn4)-positive ER tubules resist washout by both ionic and nonionic detergent after fixation, unlike non-LCV associated ER^19^. This result is particularly interesting as protein antigens localized to endomembranes are typically disrupted by both ionic and nonionic detergent permeabilization^42^, implying that Rtn4 at the LCV exhibits unique properties compared to “normal” ER-associated Rtn4. Now, with the opportunity to compare Rab5 stability at the LCV between virulent and avirulent *L.p.*, we wanted to determine whether detergent resistance was a general property of proteins recruited to the WT LCV.

Before proceeding with this experiment, we optimized a dual stain method to differentially label extracellular versus internalized bacteria, ensuring that our analysis excluded extracellular *L.p.* bound to host cells. Previous protocols relied on primary antibody labeling of bacteria while host cells were still alive^43^, allowing infection to proceed longer than desired for our application, or required multiple rounds of primary and secondary staining post-fixation^44^. However, since aldehyde fixation sufficiently permeabilizes cell membranes for primary antibodies to access cytosolic proteins to some degree^45^, we set out to improve this workflow.

We leveraged AlexaFluor dye-conjugated secondary antibodies, which are cell impermeant due to their bulk and charge at physiological pH^46^. As we opsonize *L.p.* before infection, we hypothesized that extracellular bacteria could be specifically stained post-fixation by incubating samples with an AlexaFluor conjugated secondary antibody targeting the opsonization antibody’s host species. Following permeabilization, a second round of staining with a different fluorophore-conjugated secondary antibody would yield double-labeled extracellular bacteria and single-labeled intracellular bacteria. As a proof of principle, we first validated that primary but not secondary antibodies could cross the membrane in uninfected cells post-fixation in our hands (**Fig. S3A-B**). We then infected cells, carried out the dual stain workflow, and measured mean Rab5 intensity for the bacterial cell and the local background (see Methods) for both double-labeled and single-labeled bacteria. To calculate normalized Rab5 intensity for this and all subsequent experiments, we divided mean Rab5 intensity in the *L.p.* region by the mean background and performed a log2 transformation to partially correct skew in this dataset. Normalized Rab5 intensity for single-labeled bacteria was significantly higher than for double-labeled bacteria (**Fig. S3D**). Moreover, Rab5 intensity values for double-labeled bacteria are almost entirely confined to the first three quartiles of the dataset, whereas single-labeled bacteria intensities spanned both high and low values corresponding to Rab5-positive and negative LCVs (**Fig. S3F**).

Once the dual-stain protocol was optimized for our application, we applied this workflow to cells infected with *L.p.* WT or *dotA* and permeabilized with either 0.5% saponin, a gentle surfactant that disrupts the cholesterol-rich plasma membrane but largely preserves membrane-bound epitopes^42^, or 5% sodium dodecyl sulfate (SDS), an anionic detergent that permeabilizes all endomembranes. We then performed immunofluorescence analysis of endogenous Rab5A. To confirm the efficacy of our permeabilization strategy, we first measured background-normalized total Rab5A fluorescence in uninfected cells after saponin or SDS treatment. We found that 5% SDS significantly reduced total Rab5A fluorescence, indicating successful washout of endosome membranes (**Fig. 3AB**). Next, we assessed Rab5 signal at WT and *dotA* LCVs after gentle or harsh detergent treatment. As expected, we observed many Rab5-positive LCVs for both strains in the gentle permeabilization condition. In contrast, Rab5A associated with WT LCVs resisted 5% SDS washout, whereas little to no Rab5A remained on *dotA* LCVs (**Fig. 3C**). To quantify this, we measured normalized Rab5 fluorescence at the LCV as described for the dual stain method above for each experimental condition. Across biological replicates, the mean normalized Rab5 LCV intensity was similar between the strains in the gentle permeabilization condition. In contrast, the mean normalized Rab5 LCV intensity was significantly higher in the SDS washout condition for WT *L.p.* compared to *dotA* (**Fig. 3D**), indicating that, like Rtn4, Rab5A is exceptionally stable when associated with the WT LCV.

**Figure 3:**
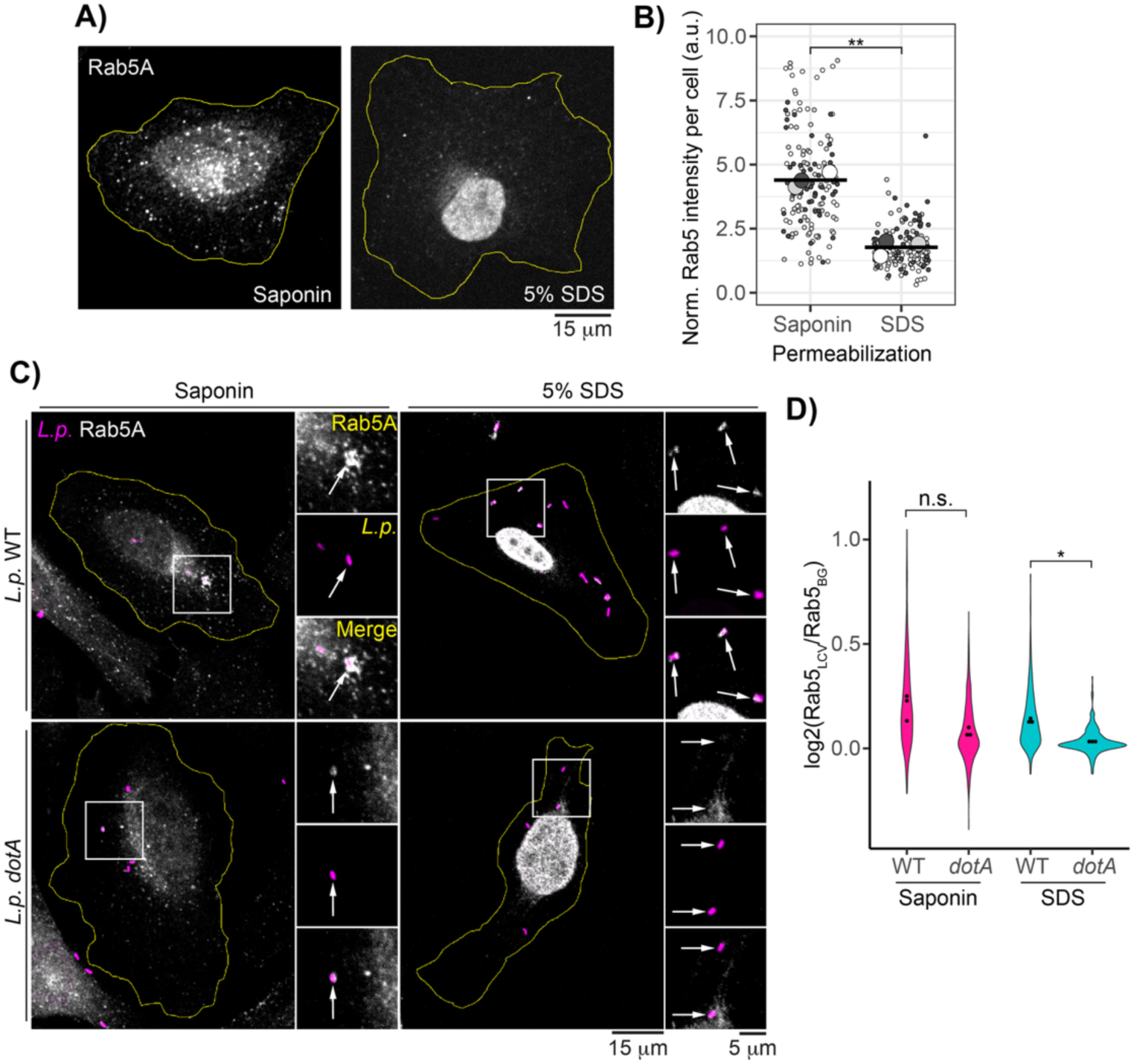
Rab5 is detergent resistant when associated with the WT LCV. (A) Saponin vs SDS permeabilized uninfected cells stained for endogenous Rab5A. (B) Quantification of normalized total Rab5 fluorescence in cells permeabilized as in panel A. N = 3, 50 cells scored per condition per replicate. Welch’s two-sample t-test, p = 0.00069.(C) Endogenous Rab5A immunofluorescence in infected cells after either saponin or SDS permeabilization. (D) Quantification of experiment described in panel C. N = 3, 80-250 LCVs scored per condition per replicate. Welch’s ANOVA followed by Games-Howell post hoc test, n.s. p>0.05, *p<0.05. For all infection experiments, cells were fixed at 1hpi.

### Morphology of ubiquitin conjugates surrounding the LCV is dependent on the SidE family of effectors

Having observed the unusual morphology and stability of Rab5 at the WT LCV, we wondered whether these characteristics were shared with other LCV-associated proteins. It is well established that polyubiquitin conjugates are recruited to the LCV starting early in infection^14,15^, with both the SidC and SidE families of effectors playing roles in the dynamics of ubiquitin recruitment. While SidC/SdcA are required for ubiquitin recruitment to the LCV at early time points post infection^27^, with a third family member, SdcB, contributing to ubiquitin recruitment in later stages^31^, the SidE family knockout strain exhibits an increase in the fraction of ubiquitin-positive LCVs during early infection ^23,47^. The phenotype is attributed to the activity of the SidE family’s DUB domain, which acts on canonically linked polyubiquitin conjugates. We reproduced this result but found that complementation of the knockout strain with plasmid encoded wildtype SdeB also recruited ubiquitin at a relatively high rate compared to *L.p.* WT, perhaps due to the overexpression of SdeB driving excess PR-ubiquitination (**Fig. 4A**). In analyzing these data, we were struck by the morphological differences in the ubiquitin-positive regions around the LCV between these strains. Around the WT LCV, ubiquitin conjugates often formed a diffuse cloud extending outwards from the bacterial cell, whereas ubiquitin signal was largely confined to the bacterial cell region for the SidE family knockout strain (**Fig. 4B**). Complementation of the SidE family knockout strain with plasmid-encoded WT SdeB rescued cloud formation, whereas complementation with the EE/AA mART mutant resulted in tight ubiquitin localization. Quantification across biological replicates showed that the ubiquitin-positive region, normalized to bacterial cell size, was significantly larger for the WT and Δ*sidE/sdeABC* + pSdeB WT strains compared to the Δ*sidE/sdeABC* and Δ*sidE/sdeABC* + pSdeB EE/AA strains (**Fig 4C**).

**Figure 4:**
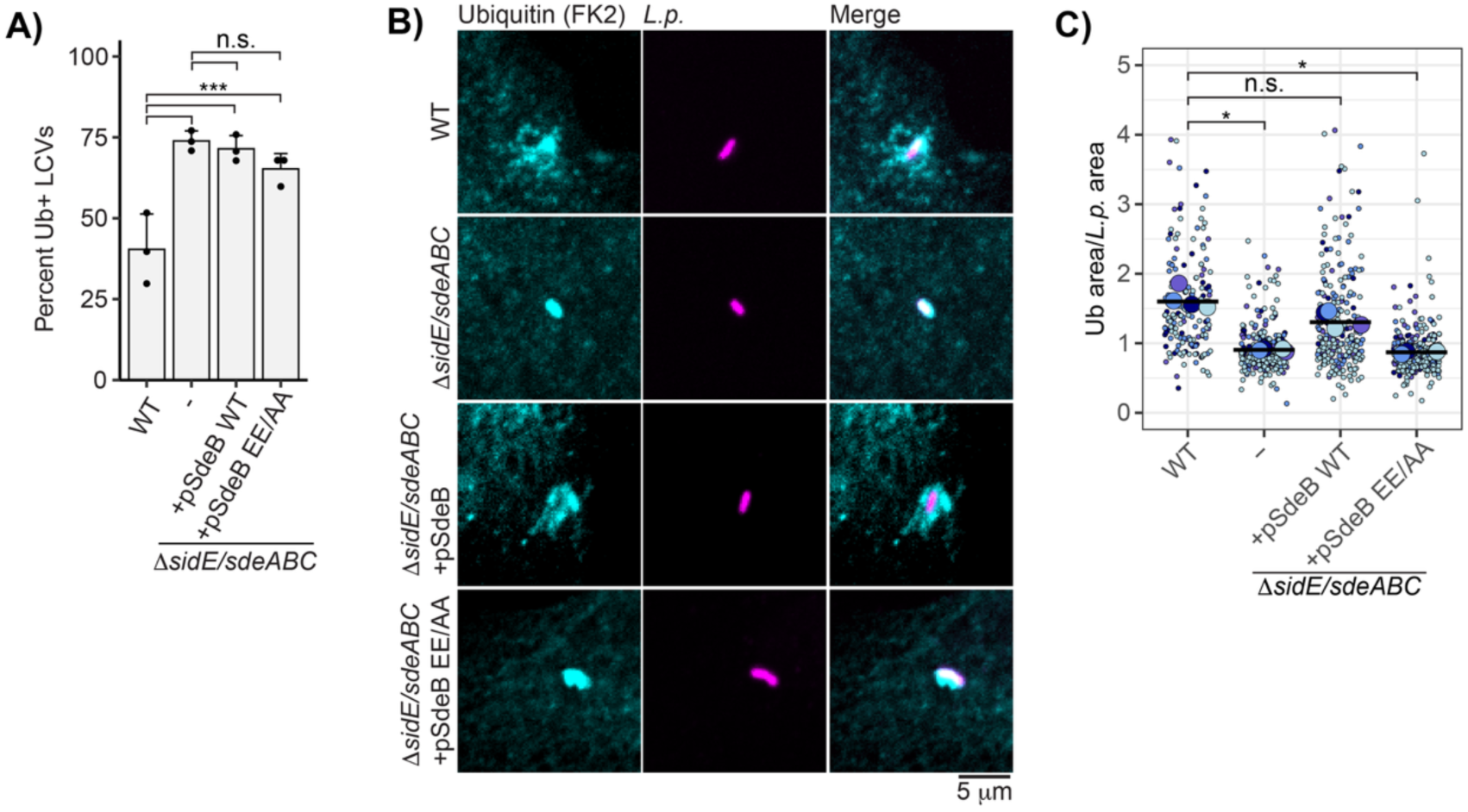
The SidE family of effectors regulates ubiquitin morphology at the LCV. (A) Manual scoring immunofluorescence imaging of endogenous ubiquitin recruitment to the indicated *L.p.* strain. N = 3, 60-120 LCVs scored per strain per replicate. G test, Bonferroni adj. p value = .01. *** p<0.0001. (B) Representative images of endogenous ubiquitin recruitment to indicated strain in experiment described in panel A. (C) Quantification of LCV-associated ubiquitin area normalized to bacterial cell area for indicated strain for experiment described in A. N = 4, 20-120 LCVs scored per strain per replicate. Welch’s ANOVA followed by Games-Howell post hoc test, n.s. p>0.05, *p<0.05.

### The SidE family of effectors is required for LCV-associated ubiquitin detergent resistance

Upon observing a similar pattern of morphological differences for LCV-localized ubiquitin between WT and Δ*sidE/sdeABC* strains, as seen previously for LCV-localized Rab5 in WT versus *dotA* strains, we next examined whether the cloud-like morphology of ubiquitin is correlated with detergent resistance. To this end, we repeated the harsh versus gentle permeabilization experiment previously described for Rab5, and quantified ubiquitin intensity at the LCV for each experimental condition. As expected, in the gentle saponin permeabilization condition many LCVs were ubiquitin-positive for all strains. While ubiquitin surrounding the WT LCV resisted SDS washout, LCVs harboring the SidE family knockout strain showed a stark reduction in ubiquitin staining after SDS treatment despite robust recruitment under the gentle permeabilization condition (**Fig. 5A**, **Fig. S4** for complemented strains). Complementation of the SidE family knockout strain with plasmid-encoded SdeB WT restored ubiquitin detergent resistance, whereas complementation with the SdeB EE/AA mutant did not (**Fig. 5B**). Next, we assessed whether the ubiquitin stabilization during *L.p.* WT infection was limited to the LCV. To do this, we measured the mean fluorescence intensity in three small cytoplasmic regions per cell in the WT-infected SDS-permeabilized condition for both infected and uninfected cells. Normalizing to extracellular background, we observed no difference between the average cytosolic ubiquitin intensity for WT-infected and uninfected cells (**Fig. 5C**), suggesting that ubiquitin stabilization predominately occurs at the surface of the LCV.

**Figure 5:**
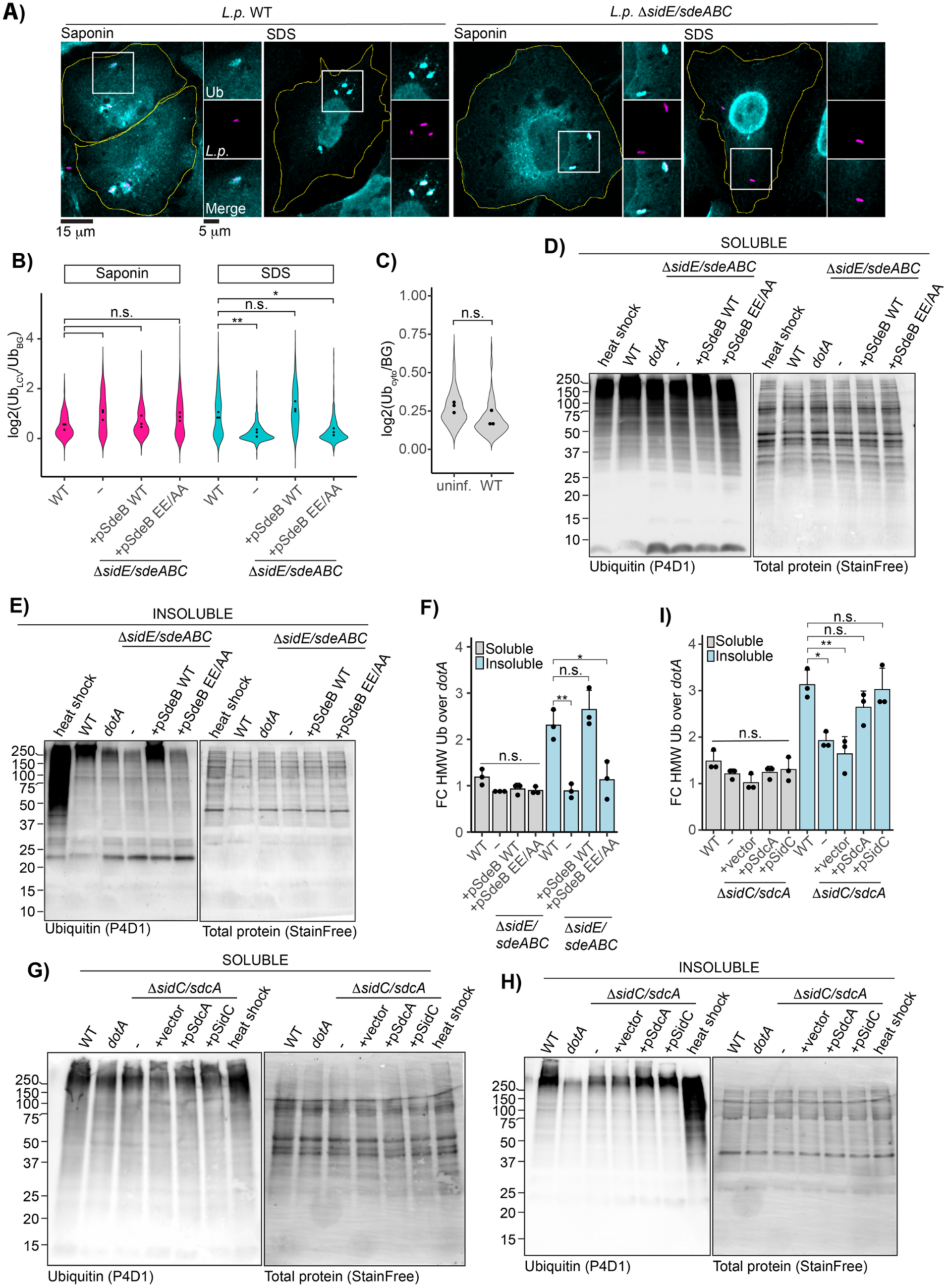
Ubiquitin at the WT LCV resists detergent washout in a SidE family dependent manner. (A) Endogenous ubiquitin immunofluorescence in cells infected with the indicated strain and permeabilized with either saponin or SDS. (B) Quantification of normalized ubiquitin fluorescence at the LCV for experiment described in A. N=3, 87-240 LCVs scored per strain per replicate. ANOVA, Tukey Kramer post hoc test, n.s. p>0.05. (C) Background-normalized cytosolic ubiquitin intensity in infected or uninfected cells after SDS washout. 17-72 cells analyzed per condition per replicate. Welch’s two-sample t-test, p = 0.09334. (D)-(E) Immunoblot analysis of total ubiquitin in the (D) soluble or (E) insoluble fraction of cells infected with the indicated *L.p.* strain from the Δ*sidE/sdeABC* panel, heat shocked, or left untreated, with total protein (StainFree, BioRad) shown as the loading control. (F) Quantification of normalized high molecular weight ubiquitin in experiments described for D and E. Total endogenous ubiquitin intensity was measured at 150 kDa and above and normalized to total protein and the fold change over the *dotA* infected control was calculated. N=3, ANOVA, Tukey Kramer post hoc test when applicable, n.s. p>0.05, * p<0.05, ** p<0.005. (G)-(I) Western blot analysis and quantification of soluble-insoluble fractionation carried out on cells infected with the *L.p.* Δ*sidC/sdcA* strain panel as described for D-F.

While the results of these immunofluorescence experiments were compelling, we sought an orthogonal approach to assay ubiquitin stability during infection. We adapted a fractionation protocol to separate soluble versus insoluble proteins to assay polyubiquitin conjugate solubility during *L.p.* infection^48^. Soluble proteins were extracted by gentle lysis in nonionic detergent, while insoluble proteins were recovered by resuspending the remaining debris in a buffer containing urea and SDS. As a positive control, we included cells subjected to heat shock, which induces the accumulation of insoluble polyubiquitinated protein aggregates^49^. After infection with *L.p.* WT, *dotA*, or the Δ*sidE/sdeABC* strain panel for one hour, levels of soluble ubiquitin conjugates were comparable across all conditions (**Fig. 5D**). On the other hand, infection with *L.p.* WT and SdeB WT complemented knockout strains increased high molecular weight insoluble ubiquitin conjugates in host cells compared to the *dotA* infected control. Insoluble ubiquitin conjugates in cells infected with the SidE family knockout and SdeB EE/AA complemented strains were equivalent to the *dotA* control (**Fig. 5E**). To quantify this effect, we focused on high molecular weight species (≥150 kDa), as WT infection appeared to specifically increase insolubility of ubiquitin conjugates in this size range. We measured integrated intensity of ubiquitin and total protein (StainFree) in this portion of each lane, normalized ubiquitin to total protein, and calculated the fold change over *dotA* control. We observed that WT infection significantly enriched insoluble ubiquitin conjugates compared to the Δ*sidE/sdeABC* strain. Complementation of the SidE family knockout strain with SdeB WT, but not EE/AA, rescued this phenotype (**Fig. 5F**). No significant differences were observed in the accumulation of high molecular weight ubiquitin conjugates between samples in the soluble fractions (**Fig. 5F**).

Finally, we repeated the solubility fractionation experiment on cells infected with the *L.p.* Δ*sidC/sdcA* strain panel. While soluble high molecular weight ubiquitin conjugates were again comparable between samples (**Fig. 5G**), infection with *L.p.* Δ*sidC/sdcA* failed to induce WT-levels of insoluble ubiquitin accumulation, a defect rescued by plasmid complementation with either SidC or SdcA (**Fig. 5H**). We note that the reduction in insoluble ubiquitin during infection with *L.p.* Δ*sidC/sdcA* is not as complete as observed for the SidE family knockout strain. This is unsurprising, as a small fraction of Δ*sidC/sdcA* are ubiquitin-positive at 1hpi^27^, perhaps due to the activity of SdcB^31^ or other ligase effectors. This finding is again consistent with ubiquitin stabilization occurring predominately at the LCV surface, and with SidC/SdcA initially acting upstream of the SidE family during infection by driving the accumulation of the first wave of polyubiquitin conjugates around the LCV.

### The hyperstable ubiquitin structure surrounding the LCV breaks down as infection progresses

We have established that LCV-associated polyubiquitin conjugates are exceptionally stable during early infection (1hpi). Next, we sought to understand whether this structure is maintained throughout infection. To this end, we carried out the gentle versus harsh permeabilization immunofluorescence assay on cells infected with *L.p.* WT for 1, 2, 4, 6, and 8 hours. In the gentle saponin permeabilization condition, many LCVs were coated in ubiquitin at all timepoints. While LCV-associated ubiquitin was robustly SDS-resistant at the early infection timepoints (1, 2, and 4hpi), few LCVs remained ubiquitin-positive at the 6- and 8-hour timepoints after SDS permeabilization, indicating that SDS resistance is lost at these late stages of infection (**Fig. 6AB**). We note that SDS resistance decreases in the same time window as LCV expansion due to bacterial replication (**Fig. 6B**).

**Figure 6 –.**
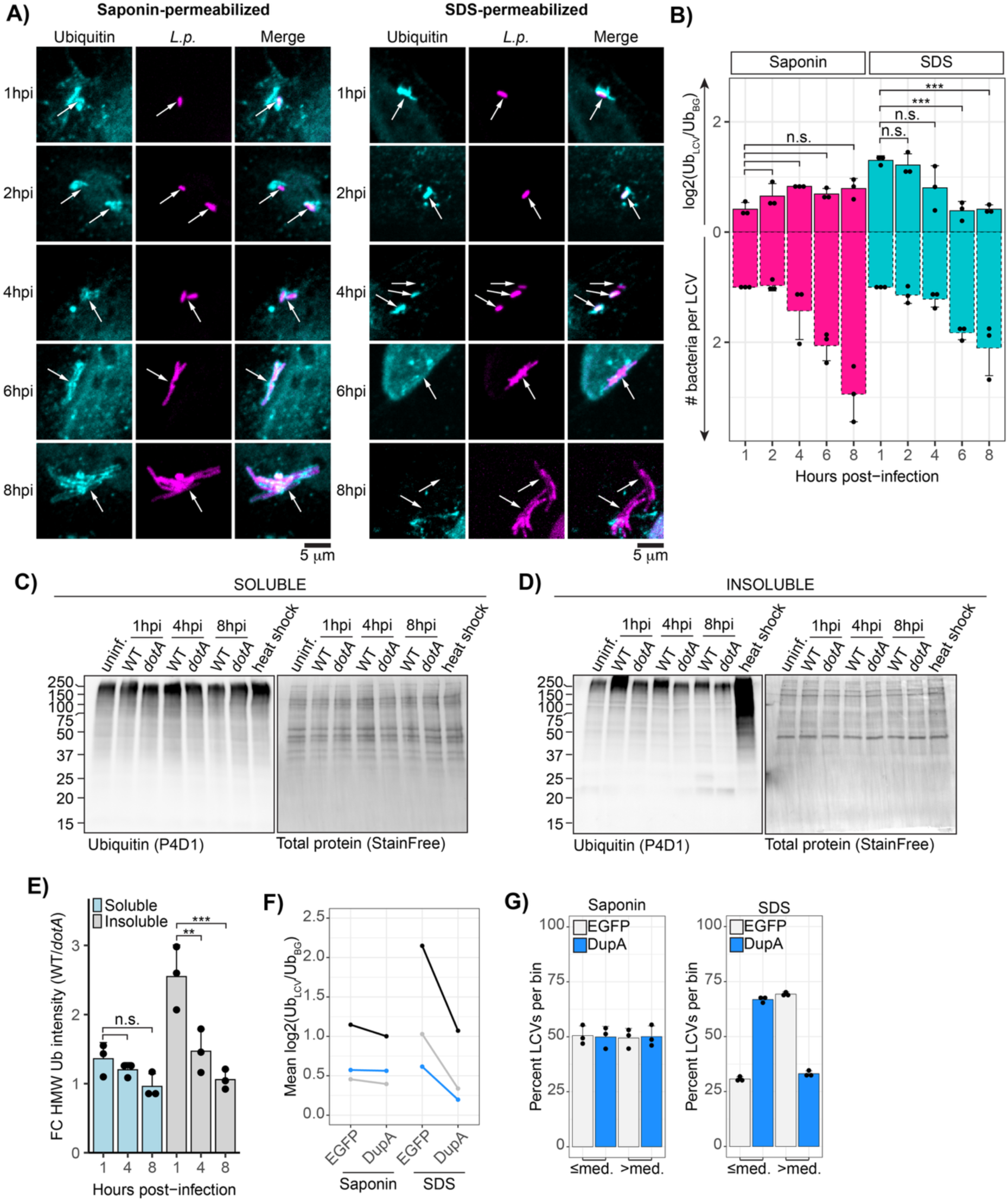
LCV-associated ubiquitin stability decreases as infection progresses. (A) Immunofluorescence analysis of endogenous ubiquitin localization to the WT LCV at the indicated timepoint post-infection after permeabilization with either saponin or SDS. (B) Normalized ubiquitin intensity quantified for the experiment described in A. Bacteria per LCV were approximated by dividing the area of each LCV by the mean LCV area for the 1hpi timepoint for each experiment and permeabilization condition. Intensity data was subjected to one way ANOVA, followed by a Tukey-Kramer post hoc test, n.s p>0.05, *** p<0.0005. (C)-(D) Immunoblot analysis of endogenous ubiquitin in cells infected with either *L.p.* WT or *dotA* for the indicated length of time and fractionated by solubility. Total protein (StainFree) is shown as a loading control. (E) Quantification of normalized high molecular weight ubiquitin in experiments described for C and E. Total endogenous ubiquitin intensity was measured at 150 kDa and above and normalized to total protein, and the fold change over the *dotA* infected control was calculated. N=3, ANOVA, Tukey Kramer post hoc test, n.s. p>0.05, ** p<0.005, *** p<0.0005. (F) Quantification of normalized ubiquitin intensity at the WT LCV at 1-hpi after saponin or SDS permeabilization in host cells expressing either EGFP or EGFP DupA. Points represent mean values for each biological replicate for a given condition, and replicates are indicated by color. (G) Analysis of the distribution of the normalized LCV-associated ubiquitin intensity values for the experiment described in panel E.

To validate the results of our immunofluorescence experiments, we fractionated cells infected with *L.p.* WT or *dotA* for 1, 4, and 8 hours by solubility. Although polyubiquitin conjugates accumulated in the insoluble fraction after 1 hour of infection with *L.p.* WT, at 4- and 8hpi we no longer observed the high molecular weight insoluble ubiquitin (**Fig. 6CD**). Quantification as described for Fig. 5 demonstrates that this finding holds true across biological replicates (**Fig. 6E**). Together, these findings suggest that the hyperstable ubiquitin structure around the LCV is a feature of early infection and breaks down as infection progresses.

### Ectopic expression of DupA destabilizes ubiquitin conjugates surrounding the LCV

Our experiments thus far suggest that SidE family-catalyzed PR-ubiquitin linkages stabilize ubiquitin around the LCV during early infection. If this is the case, cleaving PR-ubiquitin bonds in infected cells should result in a full or partial loss of detergent resistance of the ubiquitin conjugates surrounding the LCV. To test this hypothesis, we ectopically expressed DupA, an *L.p.* effector that acts as a PR-ubiquitination-specific DUB^20,50^. After verifying that our EGFP-DupA fusion construct is active against PR-ubiquitin conjugates in cells using the ubiquitin ΔGG system (**Fig. S5A**), we transfected cells with EGFP-DupA or EGFP alone, infected with *L.p.* WT for 1 hour, and performed the harsh/gentle permeabilization immunofluorescence protocol. In the gentle permeabilization condition, there was little to no difference in the mean ubiquitin intensity at the LCV between the EGFP and DupA expressing host cells. However, upon harsh permeabilization, we consistently observed a reduction in the mean ubiquitin intensity at the LCV within host cells expressing DupA compared to EGFP alone (**Fig. 6F**). Although the reduction was less pronounced than in the SidE family knockout strain—likely due to cell-to-cell variation in DupA expression—the trend was notable. While the difference between the mean intensity values between the EGFP and DupA expression conditions was not statistically significant, we felt that this data was worth investigating further. For each biological replicate, we grouped LCV intensity values by permeabilization condition and determined whether the intensity for each LCV fell above or below the median. If the intensity distributions were the same for EGFP and DupA overexpression for a given permeabilization condition, approximately 50% of the LCVs would fall above or below the median in both cases. We consistently observed this outcome for the saponin permeabilized samples (**Fig. 6G**). However, in the SDS permeabilized samples, we observed a stark shift in the distribution of intensity values: in the DupA expressing cells, 66 +/-2% of LCVs had ubiquitin intensity values below the median, compared to only 30 +/- 1% in the EGFP condition (**Fig. 6G**). Overall, this finding aligns with the hypothesis that PR-ubiquitination is essential for ubiquitin stabilization around the LCV.

### Ubiquitin expands outward from the LCV during early WT infection

We have shown that ubiquitin surrounding Δ*sidE/sdeABC* LCVs is compact and detergent sensitive during early infection, as opposed to the expansive and detergent resistant ubiquitin surrounding the WT LCV. We hypothesized that the SidE family knockout phenotype might represent an intermediate step in early ubiquitin recruitment. To explore this possibility, we used time lapse imaging to observe the morphological dynamics of LCV-associated ubiquitin during early infection. Cells were transfected with EGFP-ubiquitin and infected with *L.p.* (WT or Δ*sidE/sdeABC*) harboring a plasmid constitutively expressing unfused HaloTag and stained with the fluorescent JF646 ligand^51,52^. Imaging began within 15 minutes of infection, capturing frames every 1.5 minutes for 1-2 hours. In WT infected cells, initial ubiquitin signal often conformed tightly to the bacterial cell, then rapidly expanded outward as infection progressed (**Fig. 7A**, **movie S1**). In some cases, expansion was not observed during imaging, and in rare instances, ubiquitin recruitment appeared cloud-like from the onset (**Fig. S6AB**). This last phenotype might be explained by technical limitations, i.e. undetectably low ubiquitin signal close to the LCV before expansion. Alternatively, it might represent true biological variability in which nearby structures (such as ER membrane) are ubiquitinated by bacterial effectors and subsequently recruited to the LCV. In contrast to the WT LCVs, the vast majority of Δ*sidE/sdeABC* LCVs exhibited tight ubiquitin localization throughout the imaging time course (**Fig. 7A**, **movie S2**). Rare cases of ubiquitin expansion in the knockout strain featured highly mobile ubiquitin structures, unlike the relatively static clouds around WT LCVs (**Fig. S6C**).

**Figure 7:**
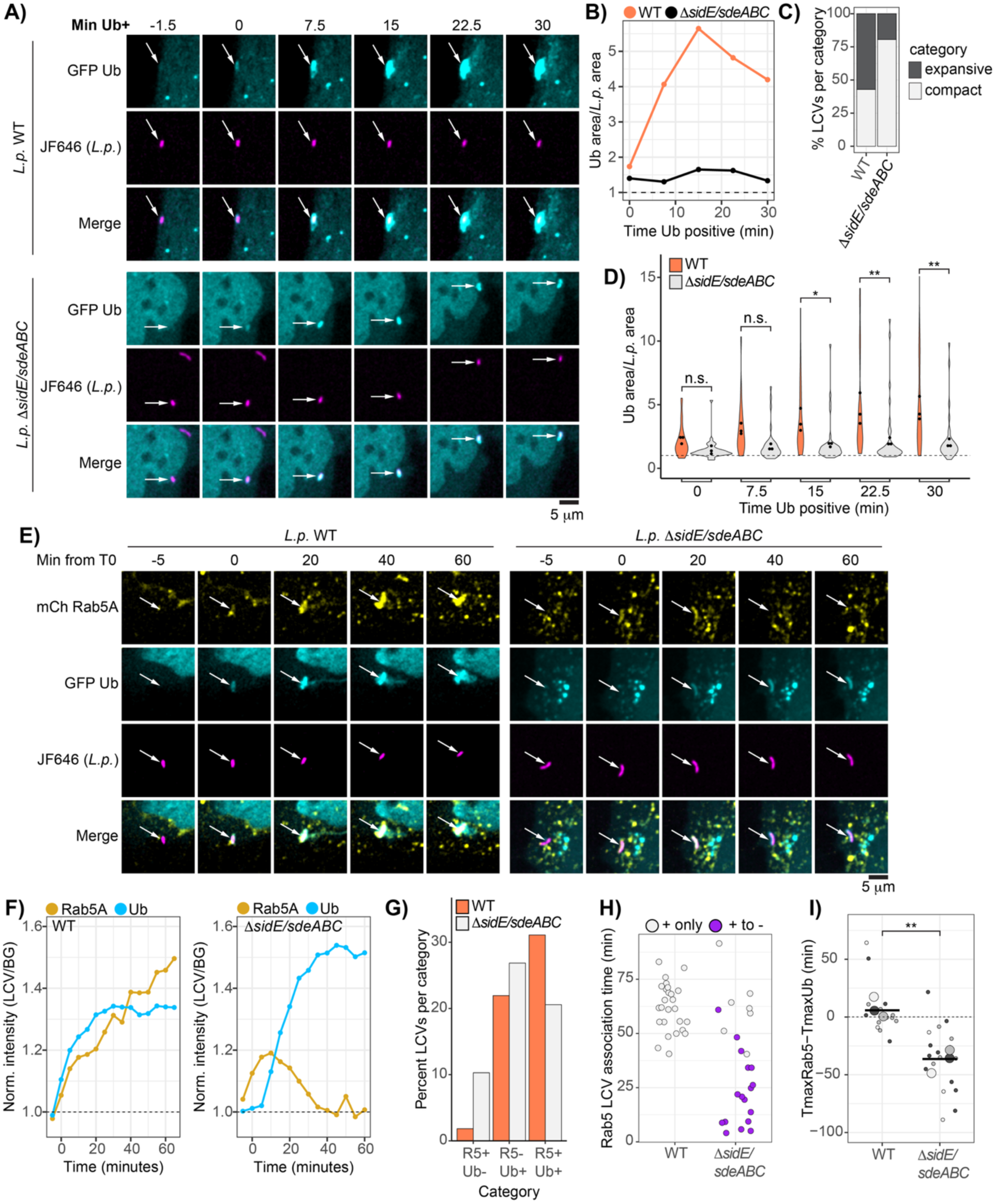
Ubiquitin expands outwards from the WT LCV during early infection and is co-recruited with Rab5. (A) Insets from live imaging time courses (see movies S1 and S2) of cells expressing EGFP ubiquitin infected with indicated *L.p.* strain labeled using the HaloTag system. (B) LCV-associated ubiquitin area normalized to bacterial cell area for LCVs shown in panel A. (C) Percentage of LCVs falling into “compact” or “expansive category. “Compact” defined as normalized ubiquitin area at T30/T0≤1.5, “expansive” indicates normalized ubiquitin area at T30/T0>1.5. Data was pooled across 3 biological replicates, 49 LCVs total for *L.p.* WT, 41 for *L.p.* Δ*sidE/sdeABC*. (D) Normalized LCV-associated ubiquitin area across imaging time course for indicated *L.p.* strains. Violin plots show distribution of pooled data, and dots indicate mean values for each biological replicate. N=3, 6-30 LCVs scored per replicate. ANOVA, Tukey Kramer post hoc test, n.s. p>0.05, *p<0.05, **p<0.005. (E)-(F) Insets from timelapse imaging in infected cells expressing mCherry Rab5 and EGFP ubiquitin, infected with either *L.p.* WT or Δ*sidE/sdeABC* (see movies S3 and S4). Graphs in F are intensity measurements for each FP tagged protein for these insets. (G) Visual scoring of ubiquitin and Rab5 recruitment patterns across all replicates. R5 is Rab5, Ub is ubiquitin. Percentages are calculated for all pooled data. 164 LCVs were score in total for *L.p.* WT, and 175 for *L.p.* Δ*sidE/sdeABC*. (H) Manual approximation of Rab5 association time with either the WT or Δ*sidE/sdeABC* LCV across 3 biological replicates. Each point represents a single LCV. LCVs that transition from Rab5 positive to negative are labeled as + to -, whereas LCVs that remain Rab5 positive at the end of the time course are marked as + only. 26 LCVs were scored for *L.p.* WT, and 24 for *L.p.* Δ*sidE/sdeABC*. (I) Comparison of the timepoints at which the intensity maximum occurs for Rab5 and ubiquitin for each LCV across three biological replicates. 3-9 LCVs were scored per strain per replicate. Welch’s two-sample t-test, p = 0.00473.

To quantify expansion of LCV-associated ubiquitin, we measured the ratio of the ubiquitin-positive region to the bacterial cell area (normalized ubiquitin area) at 7.5-minute intervals for multiple LCVs across three experiments. Only LCVs for which we observed the transition from ubiquitin-negative to positive and trackable for at least 30 minutes were included. As shown for the example insets in Fig. 7A, the normalized ubiquitin area measurement tracks with visual assessment of ubiquitin morphology dynamics for each LCV (**Fig. 7B**). To assess dataset-wide trends, we calculated the fold change in normalized ubiquitin area from T0 (first observed ubiquitin recruitment) to T30 (final timepoint analyzed) for each LCV. Applying a stringent cutoff, compact LCVs were defined as those with a fold change of 1.5 or less. We found that 80% (33/41) of the SidE family knockout LCVs fell into the compact category, compared to 43% (21/49) of WT LCVs (**Fig 7C**).

Assessing normalized ubiquitin area across the imaging time course, we found that WT LCVs displayed a steady increase from T0 to T30, whereas the SidE family knockout strain maintained a compact morphology. Notably, the mean normalized ubiquitin area at the first two timepoints quantified was comparable between the WT and SidE family knockout strains, indicating similar ubiquitin morphology upon initial recruitment. However, at later timepoints, WT LCVs exhibited significantly larger normalized ubiquitin areas, reflecting the outward expansion of ubiquitin around WT LCVs over time (**Fig. 7D**). These observations support a model wherein ubiquitin initially accumulates on the cytoplasmic face of the LCV membrane and subsequently expands outward in a SidE family-dependent manner. This expansion occurs rapidly—often visible at the next 1.5-minute interval after initial recruitment (**movie S1**). Comparing mean normalized ubiquitin between timepoints for the WT strain, we see within 22.5 minutes, the ubiquitin surrounding the WT LCV is significantly larger than at initial recruitment (p = 0.006, ANOVA, Tukey-Kramer post hoc test), whereas no significant differences are found when comparing between timepoints for the SidE family knockout strain.

### The SidE family of effectors couples Rab5 and ubiquitin association with the LCV

Our experiments thus far illustrate striking similarities in the morphology and properties of LCV-associated Rab5 and ubiquitin. Both form expansive structures in a SidE family-dependent manner and resist detergent washout when associated with the WT LCV. To assess the relative timing of ubiquitin and Rab5 LCV recruitment, as well as any subsequent morphological shifts, we leveraged our live imaging system to track these host proteins simultaneously during infection. Cells expressing EGFP-ubiquitin and mCherry-Rab5 were infected with *L.p.* WT or Δ*sidE/sdeABC* HaloTag strains as described above and imaged every 5 minutes for 1-2 hours. We focused our initial analysis on LCVs that showed both Rab5 and ubiquitin association during the imaging time course. In the *L.p.* WT case, we observed tight coordination between Rab5 and ubiquitin recruitment to the LCV (**Fig. 7EF**, **movie S3**). In contrast, Rab5 and ubiquitin LCV association was decoupled in the *L.p.* Δ*sidE/sdeABC* case, with Rab5 association most frequently preceding ubiquitin recruitment (**Fig. 7EF**, **movie S4**). Importantly, the phenotypes observed for LCV-localized endogenous ubiquitin and Rab5 morphology in our immunofluorescence experiments were reproduced in our live imaging – ubiquitin and Rab5 expanded outwards from the WT LCV and conformed tightly to the Δ*sidE/sdeABC* LCV (**Fig 7E**).

To assess data-set wide trends in Rab5 and ubiquitin LCV recruitment dynamics, we analyzed three biological replicates of each infection condition. First, we tabulated the number of LCVs that were trackable for at least 30 minutes in healthy cells that fell into four categories: double negative, Rab5-positive/ubiquitin-negative, Rab5-negative/ubiquitin-positive, and double positive (**Fig. 7G**). We observed very few WT LCVs that showed Rab5 association without ubiquitin localization. Many WT LCVs were ubiquitin-positive without detectably recruiting Rab5, and approximately 30% were double positive within the course of imaging. Interestingly, we captured many more Rab5-positive Δ*sidE/sdeABC* LCVs in our live imaging than observed in our immunofluorescence experiments. This difference is likely explained by the observation that Rab5 association with the Δ*sidE/sdeABC* LCV was transient in most cases. To quantify this phenotype, we measured Rab5 LCV association time for both strains. We included only double positive LCVs, as ubiquitin recruitment requires effector activity and Rab5-positive/ubiquitin-negative LCVs may represent failed infections. Many LCVs were Rab5-positive at the end of the imaging time course, preventing precise quantification of Rab5 association time. To address this, we included only LCVs in which Rab5 association was observed in the first half of the imaging time course, and in plotting the data differentially labeled LCVs for which Rab5 association and dissociation were observed (+ to -), as opposed to the LCVs which were Rab5-positive at the end of the time course (+ only). All WT LCVs analyzed remained Rab5-positive at the end of the time course, indicating a prolonged association time. In contrast, the bulk of the Δ*sidE/sdeABC* LCVs showed transient Rab5 association, ranging from a single imaging timepoint to approximately 60 minutes (**Fig. 7H**).

Finally, we devised a metric to assess relative timing of Rab5 and ubiquitin recruitment. After measuring normalized LCV intensity of Rab5 and ubiquitin across the imaging time course, we compared the timepoint at which the maximum intensity value (Tmax) for each marker occurred for each LCV, as we could define the maximum value more reliably that the initial recruitment timepoint. We calculated the difference between Tmax for Rab5 and ubiquitin for each LCV (11Tmax), such that a value of 0 indicates co-occurring maxima, a positive value indicates that Tmax_Ub_ precedes Tmax_Rab5_, and a negative value indicates that Tmax_Rab5_ precedes Tmax_Ub_. For *L.p.*WT, 11Tmax was close to 0 on average, indicating coupled Rab5 and ubiquitin intensity maxima. On the other hand, 11Tmax was less than 0 for all but one LCV analyzed for *L.p.* Δ*sidE/sdeABC*, indicating that maximal Rab5 association preceded maximal ubiquitin recruitment for this strain across replicates (**Fig. 7I**). These findings further support our hypothesis that incorporation of Rab5 into the proteinaceous cloud surrounding the LCV is dependent on the SidE family of effectors and demonstrate that recruitment of Rab5 and ubiquitin to the WT LCV is temporally linked.

## DISCUSSION

To survive within the host cell, intravacuolar bacterial pathogens must efficiently remodel the organelle within which they reside to support replication, evade detection, and withstand host attacks. The mechanisms contributing to each of these three missions are often inseparable, and as such gaining understanding in one realm informs all three. In this study, we explore the unique properties and formation of a hyperstable proteinaceous structure assembled rapidly around the LCV during early *L.p.* infection. Multiple host proteins, including the small GTPase Rab5A and polyubiquitin conjugates, are incorporated into this structure, adopting a distinctive “cloud” morphology around the LCV that resists harsh detergent washout. We find that the SidE family of bacterial effectors, which catalyzes non-canonical PR-ubiquitination, is required for the formation of this structure. This structure is disassembled as infection progresses on a timeline that correlates with vacuolar expansion, demonstrating that *L.p.* dynamically reshapes the properties of the proteins associated with the LCV as the bacterium’s needs shift during its intracellular life cycle.

What advantage might an expansive, stable “cloud” of protein around the LCV confer to *L.p.*? One model pertains to the vacuole guard hypothesis, the idea that intravacuolar pathogens secrete effectors that maintain the integrity of the vacuole by preventing membrane disruption ^53^.

*L.p.* effector SdhA^54,60^, and, more recently, the SidE family of effectors^32^, have been identified as vacuole guards. In the latter study, it was proposed that PR-ubiquitination of Rtn4, an ER structural protein, stabilizes ER tubules around the LCV to exclude unwanted host factors. Our experiments on Rab5 and ubiquitin suggest that many host proteins beyond Rtn4 may be incorporated into such a barrier. Further, while to our knowledge “cloud” morphology of LCV-associated proteins has heretofore not been quantified, published images of a variety of host cell types reveal expansive morphology of LCV-associated Rab6A and Rab33B^55^, Rtn4^19^, and ubiquitin^22^. In addition to building a physical barrier, the SidE family protects the LCV by camouflaging ubiquitin conjugates. Phosphoribosylation of polyubiquitin chains prevents binding of the host autophagy adaptor p62 *in vitro*^23,24^, and the SidE family knockout LCV is targeted by p62^22–24^. The SidE family of effectors appears to be a versatile player in vacuolar protection during infection, and the extent to which barrier formation and camouflage might be coupled during infection remains to be determined.

How might PR-ubiquitination induce detergent resistance and the expansive morphology we observe for host proteins around the LCV? The unique chemistry of PR-ubiquitination, in coordination with canonical ubiquitination, opens the possibility of complex intermolecular linkages between modified target proteins. A recent publication provides evidence that host proteins can be crosslinked during infection by forming PR-ubiquitin bridges between ubiquitinated substrates^24^. These crosslinks might result in a macromolecular meshwork around the LCV that could be maintained in the absence of an anchoring membrane. The expansive morphology of such a structure could be a result of disruption of molecular packing of polyubiquitin chains, and/or crosslinking to nearby cellular structures. The broader implications of this hyperstable structure are intriguing. Whether incorporation of host proteins is selective or indiscriminate remains unclear. Integration into such a crosslinked network likely impacts the activity of host proteins such as GTPases that play key roles in LCV maturation^29,56,57^. Additionally, understanding how these structures influence membrane dynamics, derived from both ER tubules and the LCV, remains an exciting avenue for future research.

How is the ubiquitin “cloud” around the LCV constructed? We provide evidence that this structure may be built in two phases: an initial wave of canonical ubiquitination, followed by co-modification by both the SidE and SidC/SdcA effector families. First, we demonstrate that the Δ*sidE/sdeABC* LCV is coated in compact, detergent-sensitive ubiquitin shortly after infection, in contrast to the expansive, detergent resistant ubiquitin surrounding the WT LCV. Using live cell imaging, we observe that ubiquitin tightly surrounds the WT LCV shortly after infection, then expands outward as infection progresses. In contrast, ubiquitin surrounding the SidE family knockout strain remains tightly LCV-associated. Further, both the SidE and SidC/SdcA families are required for maximal accumulation of detergent insoluble polyubiquitin conjugates during infection. Elegant biochemical experiments performed by Wan and colleagues^24^ demonstrated that the SidE and SidC/SdcA effector families co-modify Rab33B, resulting in mixed canonical and PR-linked ubiquitin chains. Our data is consistent with a model in which canonically linked ubiquitin accumulates around the LCV in a SidC/SdcA dependent manner shortly after infection, at which point the ubiquitin coat is detergent-sensitive and compact. Subsequently, PR modification and crosslinking by the SidE family of effectors, as well as further canonical ubiquitination by SidC/SdcA as suggested by Wan et al, generates the expansive, detergent resistant “cloud” of polyubiquitin conjugates around the LCV. While we suspect that, in the presence of both the SidE and SidC/SdcA effector families, substrate proteins are rapidly co-modified upon LCV association, the accumulation of ubiquitin around LCVs harboring *L.p*. Δ*sidE/sdeABC*, but not *L.p*. Δ*sidC/sdcA*^27,31^, points to the essentiality of canonical ubiquitination driven by SidC/SdcA in initiating construction of the hyperstable polyubiquitin structure during infection. We emphasize that we do not argue that canonical ubiquitination must proceed PR-ubiquitination biochemically, but rather that, during infection, the activity of canonical ubiquitin ligase effectors is required for initial ubiquitin accumulation around the LCV, providing a properly localized substrate for SidE family modification. The role of additional regulatory mechanisms in constructing the ubiquitin cloud, such as the SidE family’s N-terminal DUB domain implicated in ubiquitin^47^ and ER tubule^23^ dynamics at the LCV surface, remains an open question.

The complex interplay between ubiquitination and LCV membrane association is illustrated by the phenotypic differences in Rab5 recruitment relative to ubiquitin accumulation around the LCV for *L.p.* WT and Δ*sidE/sdeABC*. While LCV recruitment of Rab5 and ubiquitin appears tightly coordinated in the *L.p.* WT case, Rab5 association with the Δ*sidE/sdeABC* LCV, when captured in our imaging, is transient and precedes ubiquitin accumulation. Previously, our lab demonstrated that Rab5 is mono-ubiquitinated during infection with the *ΔsidE/sdeABC* strain without the accumulation of exceptionally high molecular weight species^28^. While Rab5 ubiquitination is largely dependent on SidC/SdcA during infection^28,30^, a recent report provides evidence that another effector, lpg2525, can also ubiquitinate Rab5, and that in the absence of this effector Rab5 accumulates at the LCV surface at high levels^58^. Intriguingly, we observe the opposite phenotype with the Δ*sidC/sdcA* strain, in which loss of Rab5 ubiquitination correlates with loss of LCV association^28^. This apparent conflict may be explained by tight coordination between SidC/SdcA and the SidE family, in which co-modification by both effector families locks Rab5 into place at the LCV surface, as previously discussed. Further, we have not eliminated the possibility that SidC/SdcA recruits Rab5 to the LCV through means other than direct ubiquitination. Taken together, we hypothesize that, in the Δ*sidE/sdeABC* infection case, Rab5 is recruited to the LCV via the activity of either host or bacterial proteins, and then canonically ubiquitinated, which, in the absence of the SidE family, drives dissociation from the membrane. Why Rab5 recruitment consistently precedes ubiquitin accumulation at the Δ*sidE/sdeABC* LCV remains an open question, and further investigation will likely yield insights into the intricate web of bacterial effectors regulating the relationship between small GTPase LCV association and ubiquitination.

While Rab5 recruitment to endocytic vesicles within eukaryotic cells usually confers “early endosome”-like properties, data from our lab and others indicates that *L.p.* effectors divert the vacuole from “normal” progression through the endolysosomal system. Previously, we found that the Rab5 binding partner EEA1, required for homotypic fusion, is excluded from the WT LCV membrane in the same time window as peak Rab5 recruitment^28^, and it has been demonstrated that the WT LCV resists fusion with early endosomes^59,60^. Further, it is well established that *L.p.* effectors remodel the phosphoinositide composition of the vacuole membrane from endosome-like (PI(3)P+ or PI(3,5)P2+) to predominately PI(4)P+ during early infection^9,61,62^. Interestingly, the Rab5 GEF Rabex5 binds ubiquitin; this is thought to be one mechanism by which Rab5 is targeted to membranes^63^. It is possible that Rabex5 binding to ubiquitin contributes to Rab5 LCV recruitment, although if this were the case, one might predict Rabex5 overexpression to increase Rab5 recruitment to the WT LCV, which we do not observe, and for Rab5 association with the LCV to trail behind ubiquitin recruitment. Instead, we found that ubiquitin and Rab5 were recruited simultaneously to the WT LCV, and that Rab5 recruitment preceded ubiquitin association with the Δ*sidE/sdeABC* LCV. Finally, it has been reported that late endosomal GTPase Rab7 is recruited to the LCV^64^, and that this recruitment may be driven by SUMOylation^65^. While there is evidence that the WT LCV has characteristics of late endosomes/lysosomes in the end stages of infection^66^, the WT LCV resists fusion with lysosomes throughout early infection^12,33^. Our data is not consistent with a model in which the LCV undergoes classical endosome maturation from Rab5+ to Rab7+ driven by host regulators; while it is fascinating that both host GTPases are recruited to the WT LCV, this association seems to be regulated by post-translational modification of these proteins by bacterial effectors rather than the activity of host regulators.

Ubiquitin tagging of invading bacterial cells or vacuoles often corresponds to targeting by host autophagy machinery and ultimate trafficking to the lysosome, and accordingly many pathogens evade ubiquitination as part of their virulence programs^67^. It appears that *L.p.* has taken an entirely different approach and integrated ubiquitin recruitment into the maturation of its vacuole. As proposed here and elsewhere^23,24,32^, ubiquitin may serve as a glue that holds together a physical barrier around the LCV to protect the membrane from unwanted vesicular fusion or targeting by autophagy. Ubiquitin localization may also function as a molecular timer, coordinating stages of LCV maturation; a recent paper links late-stage LCV expansion to a bacterial effector phospholipase that is activated by ubiquitin^68^. Our finding that ubiquitin hyperstability at the LCV is lost as infection progresses further suggests that ubiquitin dynamics are tightly coordinated with stages of the *L.p.* life cycle within the host cell. Considering the astounding evolutionary breadth of eukaryotic cells that support *L.p.* replication^5^, embracing rather than escaping ubiquitin tagging of the LCV may allow *L.p.* to rapidly protect and remodel its vacuole in a wide variety of cell types, as ubiquitin is extremely highly conserved across eukaryotes^69^. In conclusion, our study highlights a novel role for ubiquitin in *L.p.* pathogenesis. By leveraging both canonical and noncanonical ubiquitination, *L.p.* constructs a hyperstable structure surrounding the LCV that may protect against host defenses. These findings deepen our understanding of ubiquitin’s versatility and suggest that further exploration of *L.p.*’s co-opting of host signaling pathways will reveal fundamental insights into host-pathogen interactions.

## ACKNOWLEDGMENTS

We thank Tom Moss and Dr. Joe Henry Steinbach for feedback on the manuscript. We thank Dr. Varun Bhadkamkar for thoughtful discussions of the models proposed in this study. A.S. acknowledges support from the MPHD T32 training grant and the Hooper Graduate Student Fellowship. S.M. acknowledges financial support from the National Institutes of Health (grant nos. R01GM140440 and R01GM144378), the Pew Charitable Trust (grant no. A129837), a Bowes Biomedical Investigator award, and a gift fund from the Chan-Zuckerberg Biohub.

## KEY RESOURCES

### Plasmids

| Description | ID | Source |
| --- | --- | --- |
| EGFP in pcDNA3.1 | pAS079 | This study |
| EGFP hRabex5 in pcDNA3.1 | pAS061 | This study |
| EGFP hRabapatin5 in pcDNA3.1 | pAS062 | This study |
| SdeB in pJB1806 | pAS083 | Steinbach and Bhadkamkar et al, 2024<sup>28</sup> |
| SdeB EE/AA in pJB1806 | pCM003 | This study |
| Myc-SdeA in pMSCV |  | Gift from Dr. Ralph Isberg |
| Myc-SdeB in pMSCV |  | Gift from Dr. Ralph Isberg |
| Myc-SdeC in pMSCV |  | Gift from Dr. Ralph Isberg |
| Myc-SidE in pMSCV |  | Gift from Dr. Ralph Isberg |
| Myc-LotC in pMSCV |  | Gift from Dr. Ralph Isberg |
| pcDNA3.1 | pSM022 | Gift from Dr. Craig Roy |
| 3X Flag hRab5A in pcDNA3.1 | pAS034 | Steinbach and Bhadkamkar et al, 2024<sup>28</sup> |
| HA ubiquitin in pRK5 | pSM099 | Gift from Dr. Kohei Arasaki |
| HA ubiquitin ΔGG in pRK5 | pAS117 | This study |
| DupA in pEGFPc2 | pCM004 | This study |
| Ubiquitin in pEGFPc1 | 11928 (Addgene) | Gift from Dr. Nico Dantuma |
| HaloTag in pON | pAS100 | This study |
| 3X Flag hRab5A Q79L in pcDNA3.1 | pAS039 | This study |
| 3X Flag hRab5A S34N in pcDNA3.1 | pAS040 | This study |
| 3X Flag hRab5A 1-211 in pcDNA3.1 | pAS041 | Steinbach and Bhadkamkar et al, 2024<sup>28</sup> |
| mCherry hRab5A in pcDNA3.1 | pAS042 | Steinbach and Bhadkamkar et al, 2024<sup>28</sup> |
| mCherry Rab5A S34N in pcDNA3.1 | pAS048 | This study |
| mCherry Rab5A Q79L in pcDNA3.1 | pAS047 | This study |

### Legionella strains

| Description | ID | Source |
| --- | --- | --- |
| *Legionella pneumophila* serogroup 1 strain Lp02 *rpsL hsdR thyA* ("WT") | LEG003 | Berger and Isberg, 1993<sup>11</sup>. Gift from Dr. Craig Roy |
| Lp02 *dotA* | LEG004 | Berger and Isberg, 1993<sup>11</sup>. Gift from Dr. Craig Roy |
| Lp02 $\Delta sidE$ $\Delta sdeC$ $\Delta sdeBA$ ($\Delta lpg0234$, $\Delta lpg2153$ $\Delta lpg2156-2157$), annotated as "$\Delta sidE/sdeABC$" for brevity | LEG151/JV6113 | Jeong *et al.*, 2015<sup>70</sup>. Gift from Dr. Ralph Isberg |
| Lp02 $\Delta sidE/sdeABC$ pJB1806::SdeB | LEG171 | Steinbach and Bhadkamkar *et al.*, 2024<sup>28</sup> |
| Lp02 $\Delta sidE/sdeABC$ pJB1806::SdeB EE/AA | LEG196 | This study |
| Lp02 pON::Halotag | LEG195 | This study |
| Lp02 $\Delta sidE/sdeABC$ pON::HaloTag | LEG160 | This study |

### Antibodies

| Antigen (antibody clone) | Application (dilution) | Supplier (catalog #) |
| --- | --- | --- |
| Rab5A (E6N8S) | IF (1:100) | Cell Signaling Technology (46449) |
| Flag (M2) | WB (1:2500), IP (1:50) | Sigma (F1804) |
| HA-HRP | WB (1:1000) | Thermo (26183-HRP) |
| *L. pneumophila* | Opsonization (1:2000) | Thermo (PA1-7227) |
| Myc | WB (1:1000) | Proteintech (60003-2-Ig) |
| GFP | WB (1:1000) | Roche (11814460001) |
| Ubiquitin (P4D1) | WB (1:500) | Santa Cruz (sc-8017) |
| Ubiquitin conjugates (FK2) | IF (1:200) | Millipore (ST1200) |

### Software and packages used (all code is available upon request)

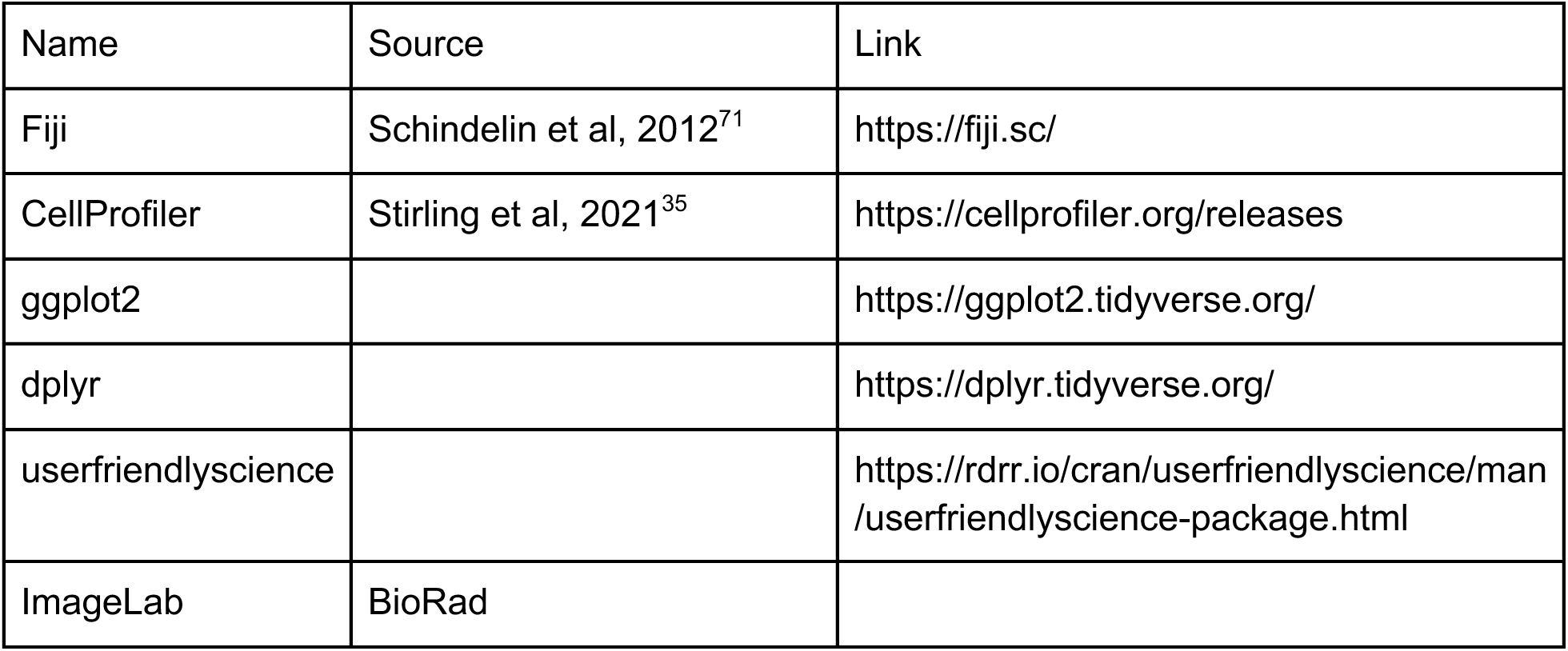

## METHODS

### Cell lines

Parental HEK293T were obtained from ATCC. The monoclonal HEK293T FcγRIIb cell line used for Western blotting was produced by lentiviral transduction, hygromycin selection, and limiting dilution using standard protocols. HeLa FcγR cells used for imaging were a gift from the lab of Dr. Craig Roy. All cell lines were regularly tested for mycoplasma contamination. Cells were maintained in Dulbecco’s modified Eagle’s medium (DMEM, Gibco) supplemented with 10% heat-inactivated fetal bovine serum (FBS, VWR) and 2 mM L-glutamine in a humidified incubator at 37°C, 5% CO_2_.

### Bacterial strains and plasmid construction

*L.p.* strains used in this study were derived from the Philadelphia-1 thymidine auxotroph stains LP02 and LP03^11^. The Δ*sidE/sdeABC* strain (JV6113)^70^ was a gift from Dr. Ralph Isberg. Plasmids were introduced into *L.p.* strains by electroporation as previously described^11^. *L.p.* genes were amplified from purified gDNA, mutagenized by PCR amplification if applicable, and inserted into the pJB1806 or pEGFPc2 vector by Gibson assembly. The constitutive expression HaloTag plasmid was derived from the mCherry pON plasmid described previously^52^. Rabex5 and Rabaptin5 genes were PCR amplified from HeLa cDNA, N-terminally fused to EGFP by overlap extension PCR, and cloned into the pcDNA3.1 expression vector. Rab5 mutant constructs were generated by overlap extension PCR of the original WT constructs using mutagenesis primers, then cloned into the pcDNA3.1 expression vector.

### Transfection of mammalian cells

Cells were transfected with high quality maxiprepped plasmid DNA (Sigma Genelute HP kit) using the jetPRIME transfection system (PolyPlus). Cells were grown to ∼70% confluence before transfection. For HEK293T FcγR, total μg of DNA was added as recommended by the supplier for the culture vessel; for HeLa FcγR, the amount of DNA was halved to reduce cytotoxicity. Cells were transfected 24 hours before further experimental procedures or harvesting.

### *L.p.* culture and infection of FcγR cell lines

Heavy patches of *L.p.* streaked from single colonies were grown on N-(2-acetamido)-2-aminoethanesulfonic acid (ACES, Sigma) buffered charcoal yeast extract plates supplemented with thymidine and, if applicable, 5 μg/mL chloramphenicol at 37°C. Heavy patches were resuspended in ACES buffered yeast extract broth (AYE) supplemented with thymidine, L-cysteine, iron nitrate, and chloramphenicol if required, and either used to start overnight liquid cultures, or used directly for infection. Overnight cultures were grown to an optical density of 3-4. When required, expression from the pJB1806 plasmid was induced with 1 mM isopropyl β-d-1-thiogalactopyranoside (IPTG) at 37°C for two hours before infection. Cultures were enumerated and the required volume for the desired multiplicity of infection (1 for imaging experiments, 20 for Western blotting) was added to a 1:2000 dilution of opsonization antibody in mammalian cell culture media. Opsonization proceeded for 20 minutes at 37°C with end-over-end mixing. To infect, opsonized bacteria were added to mammalian cells, and the culture vessels were spun for 5 minutes at 1000xg to synchronize the infection (“spinfection”). For infections longer than 1 hour, infection media was aspirated and cells washed 3X with pre-warmed 1X PBS at 1hpi. The final wash was replaced with fresh pre-warmed cell culture media, and the infection was allowed to continue in the incubator for the desired duration.

### Immunofluorescence sample preparation

HeLa FcγR cells were plated on poly-L-lysine coated glass #1.5 coverslips and grown to ∼70% confluence. Once experimental manipulations were complete, cells were washed three times in 1X phosphate buffered saline (PBS) and fixed in 4% paraformaldehyde (PFA) in PBS for 15 minutes at room temperature. After removal of PFA, coverslips were washed three times in 1X PBS. If applicable, extracellular bacteria were stained with 1:1000 goat anti-rabbit AlexaFluor 405 (Invitrogen) in 2% w/v bovine serum albumin (BSA)/0.02% NaN_3_/PBS for 30 minutes at room temperature (RT), then washed three times in 1X PBS. Cells were permeabilized either in 0.5% saponin/2% BSA/0.02% NaN3/PBS (block/perm buffer), or in 5% SDS/Tris buffered saline (K+ free) for 1 hour at RT with orbital agitation at 150 rpm. For SDS permeabilized cells, coverslips were washed in 1X PBS to remove residual detergent, then incubated for 1 hour in block/perm buffer at RT. All samples were stained overnight in primary antibody diluted in block/perm buffer at 4°C in a humidified chamber. The next day, coverslips were washed 3X in 1X PBS, then stained with secondary antibody(ies) diluted in block/perm buffer for 1 hour at RT. After removal of the secondary stain, coverslips were washed 3X in 1X PBS, then 3X in ultrapure water, briefly dried, and mounted on glass slides with Prolong Glass hard curing mountant (Thermo). Slides were cured lying flat in the dark overnight at RT, and, if required, subsequently stored at 4°C until imaging was completed.

### Halo staining of *L.p.* and sample preparation for live imaging experiments

HeLa FcγR cells were plated in 35 mm glass bottom dishes (#1.5, Cellvis) and grown to ∼70% confluence. Cells were transfected as described above. *L.p.* harboring the HaloTag pON plasmid were grown from a single colony as a heavy patch for 48 hours on CYE + chloramphenicol and suspended in AYE. 2x10^8^ bacterial cells were transferred to a sterile microcentrifuge tube and pelleted at 11,000xg for 1 minute. The supernatant was removed and the pellet was resuspended in 50 uL 5 μM JF646 (Lavis lab, Janelia). Bacteria were stained for 20 minutes at RT in the dark. After staining, bacteria were opsonized in the dark as described above. HeLa FcγR cells were spinfected as described above, then immediately washed 3X in prewarmed 1X PBS. The final wash was replaced with prewarmed DMEM without phenol red (Gibco) supplemented with heat-inactivated 10% FBS (VWR) + 2 mM L-glutamine (Sigma). Cells were immediately transferred to a humidified live imaging chamber stage insert (37°C, 5% CO_2_) (Okolabs). Imaging was initiated with 15 minutes of spinfection.

### Image acquisition

All images were acquired on a Nikon Ti2E inverted microscope with a CREST-XLight LFOV spinning disk system and Photometrics Prime95B camera (16 bit, 1x1 pixel binning) using a 60X 1.4 NA Plan Apo oil immersion lens objective. All immunofluorescence slides were blinded before imaging and analysis. NIS Elements Advanced Research software was used to control the microscope and acquire images. For fixed cells, Z stacks were acquired in 0.3 um increments across a Z range of 3 um. Exposure time ranged from 10-50 ms, with the following laser power settings: ExW 365 nm 25%, ExW 488 nm 25%, ExW 561 nm 100%, ExW 640 nm 100%. For live samples, Z stacks were acquired in 0.5 um increments across a Z range of 2 um. Laser power was reduced to 10-15%, 10 ms exposure. 8-10 XY positions were imaged every 1.5 minutes for ubiquitin only imaging, or 5 minutes for ubiquitin and Rab5 dual imaging, for 1-2 hours.

### Image analysis

Files were initially processed in Fiji. All analysis was conducted on maximum intensity Z projections. For all image analysis, only cells with 10 or fewer internalized bacteria (if applicable) that showed no signs of impending cell death (rounding, blebbing) were included. For area analysis of LCV-associated Rab5 or ubiquitin in fixed cells, LCVs staining positive for a marker of interest were selected by eye, and a small region of interest (ROI) around the positive LCV(s) was duplicated and saved as a new file. Using custom ImageJ macros, these images were then background subtracted (rolling ball radius = 10) and split into individual channels (*L.p.* and marker of interest). These images were used as the input for CellProfiler pipeline analysis. Briefly, bacterial cells were defined as primary objects, and the LCV associated marker of interest was defined as a secondary object by propagation. These objects were then related, and the area of each object was measured. For each biological replicate, all normalized area measurements were pooled, and the top 0.5% of values discarded to remove extreme outliers due to CellProfiler secondary object propagation errors. For area analysis in live cells, area for bacteria and LCV-associated ubiquitin was measured by hand in Fiji at indicated time points. For intensity analysis in fixed cells, ROIs for all intracellular bacteria were selected as above in Fiji, and images were split into individual channels (background subtracted only from the *L.p.* channel). These images were used as input for CellProfiler analysis. Bacterial cells were defined as primary objects, then expanded by 7 pixels. The expanded LCV object was then used as an inverted mask for the marker channel. Intensity was measured in the marker channel within the original bacterial cell object (LCV intensity) and in the marker channel masked with the expanded bacterial cell object (background intensity). For intensity analysis of live cell imaging, LCV and adjacent background regions were measured by hand in Fiji, and the mean LCV intensity was normalized to the mean background intensity. For endosome size analysis in uninfected cells in CellProfiler, the cell bodies of transfected cells were identified using the GFP channel after log transformation, and used as a mask for the Rab5 channel, in which endosomes were defined as primary objects and measured. For total cell fluorescence analysis in uninfected cells, cell area was defined manually in Fiji. Integrated density for cell area was measured, as well as mean intensity for three background regions for each image. For live imaging, supplemental movies and associated insets were smoothed for representation purposes using the built-in Fiji “Smooth” function (3x3 pixel averaging filter).

### Whole cell lysate preparation and Western blotting

Cells were grown in 6 well plates to ∼70-90% confluence. Once experimental manipulations were complete, media was aspirated and cells were washed 3X with ice cold 1X PBS on ice. Cells were gently scraped into 1 mL ice cold 1X PBS and transferred to microcentrifuge tubes. Cells were pelleted at 3000xg for 10 minutes at 4°C, and the supernatant was aspirated. Pellets were gently resuspended in lysis buffer (150 mM NaCl, 50 mM Tris pH 7.4, 1% Triton-X 100, plus 1 mM PMSF, 1X Roche cOmplete protease inhibitor cocktail, and 10 mM N-ethylmaleimide [NEM]). Cells were agitated for 20 minutes then centrifuged at 16,000xg for 20 minutes at 4°C. Cleared lysates were transferred to a new tube, and protein concentrations were quantified using the Pierce 660 nm Assay Kit. For each sample, 30-50 μg total protein was denatured in 1X SDS sample buffer, 1% v/v β-mercaptoethanol (BME) for 5 minutes at 95°C. Lysates were loaded onto polyacrylamide gels (10, 12, or 15%) and separated by SDS-PAGE. Proteins were transferred to 0.2 um pore PVDF membranes overnight at 4°C in 1X CAPS, 10% methanol. Membranes were blocked in 5% non-fat dry milk (BioRad) in PBS-T (0.1% Tween-20 in PBS) for 1 hour at RT. Unconjugated primary antibodies were diluted in 2% BSA, 0.02% NaN3 in PBS-T, whereas HRP conjugates were diluted in blocking buffer (see table for dilutions). Membranes were incubated with primary antibody solutions overnight at 4°C. Membranes were washed 3X in PBS-T, then incubated with 1:5000 secondary antibody in blocking buffer for 1 hour at RT. Membranes were washed 3X in PBS-T, then developed for 1 minute in Amersham ECL Western Blotting Detection Reagent (Global Life Science Solutions) and imaged on a ChemiDoc Imaging System (BioRad).

### Immunoprecipitation

Cells were grown in 10 cm culture dishes to ∼70-90% confluence. Once experimental manipulations were complete, cells were washed and pelleted as described above. Cells were lysed in ice cold IP lysis buffer (137 mM NaCl, 20 mM Tris pH 8, 1% NP40, 2 mM EDTA, plus 1X Roche cOmplete protease inhibitor, 1 mM PMSF, 10 mM NEM) and the lysate concentration was quantified as described above. Samples were diluted to equal concentrations, and input samples were removed and prepared for SDS-PAGE as described above. 300 uL of remaining sample (1-3 mg/mL) was used for immunoprecipitation with Flag-M2 conjugated magnetic beads (Millipore Sigma) following the manufacturer protocol. Immunoprecipitated proteins were eluted by boiling at 95°C in 2X SDS sample buffer, 1% v/v BME. For SDS-PAGE separation, 3-5% input and the full eluate volume for immunoprecipitated samples were loaded.

### Soluble/insoluble fractionation

Protocol was adapted from Rai et al, 2021^48^. Briefly, cells were grown in 6 well plates to ∼70-90% confluence. Heat shocked cells were incubated at 43°C, 5% CO_2_ in a humidified tissue culture incubator for 1 hour. Once experimental manipulations were complete, cells were washed and pelleted as described above for whole cell lysate preparation. Cells were resuspended by gentle pipetting in ice cold Triton buffer (1% Triton X 100 in PBS with 1 mM PMSF, 1X Roche Complete protease inhibitor cocktail, and 10 mM NEM). Cellular debris was pelleted at 16000xg for 20 minutes at 4°C, and the supernatant (soluble fraction) was transferred to a new tube. The pellet was washed 3X in ice cold Triton buffer. The insoluble debris was suspended in 100 u/mL Benzonase (MilliporeSigma), 7 M urea in RIPA buffer (Cell Signaling Technology), supplemented with 10 mM MgCl_2_ to saturate EDTA. Digestion proceeded at Rt for 20 minutes, at which point 10% SDS in ultrapure water was added to a final concentration of 1%. Samples were then agitated for 5 minutes and centrifuged at 16,000xg for 10 minutes at 4°C. The supernatant (insoluble fraction) was transferred to a new tube. Protein concentration was quantified for the soluble protein fraction as described above. Laemmli buffer plus 6% BME was added to 1X and samples were boiled for 5 minutes at 95°C. For soluble fractions, 50 μg total protein was loaded, and an equal volume was loaded for the corresponding insoluble fraction. Samples were run on 8-16% gradient StainFree gels (BioRad), which were then activated according to manufacturer instructions, transferred to PVDF, and immunoblotted as described above.

### Western blot quantification

Blots for soluble/insoluble fractionation experiments were quantified using the BioRad ImageLab software. Lanes were first defined in the desired molecular weight range using the StainFree image, then applied to the appropriate chemiluminescence image. Whole lane volume was measured for both StainFree and chemiluminescence images. Chemiluminescence signal was first normalized to StainFree signal, then to the appropriate untreated control sample to find intensity fold change as plotted in Figure 5.

### Data representation and statistics

All representative images are max intensity Z projections including only intracellular bacteria (if applicable) unless indicated. For all jitter plots, color corresponds to biological replicate. Small dots are individual measurements (e.g. normalized area of LCV-associated Rab5 for one bacterial cell), and larger dots represent the mean of a given biological replicate. Black bars represent the mean of data pooled across biological replicates. For bar plots, points are measurements for each biological replicate, bars represent the mean value across biological replicates, and error bars show standard deviation. For violin plots, points represent mean values for each biological replicate. Datasets with one nominal variable with two values were subjected to a Welch’s t-test on mean values for each biological replicate using a p-value of 0.05 as a significance threshold. For datasets with one nominal variable with more than two values, homogeneity of variance was first tested using a Bartlett test on the mean values for each biological replicate. If the Bartlett test resulted in a p value greater than 0.05, equal variance was assumed, and the data was subjected to a one-way ANOVA followed by a Tukey-Kramer post hoc test (confidence level 95%) if the ANOVA indicated significant differences between groups (p<0.01). If the Bartlett test resulted in a p value<0.05, equal variance could not be assumed, and data was subjected to a Welch’s ANOVA followed by a Games-Howell post hoc test (code from userfriendlyscience R package) if significant differences between groups were observed. For percentage data, pooled counts were subjected to a G test, followed by pairwise comparisons if significant differences were detected. Significance for pairwise comparisons was assessed using a Bonferonni-adjusted p-value (values indicated in figure captions). All data processing, statistical analysis, and plotting was done in R (dplyr and ggplot2 packages).

---

**Figure S1.**
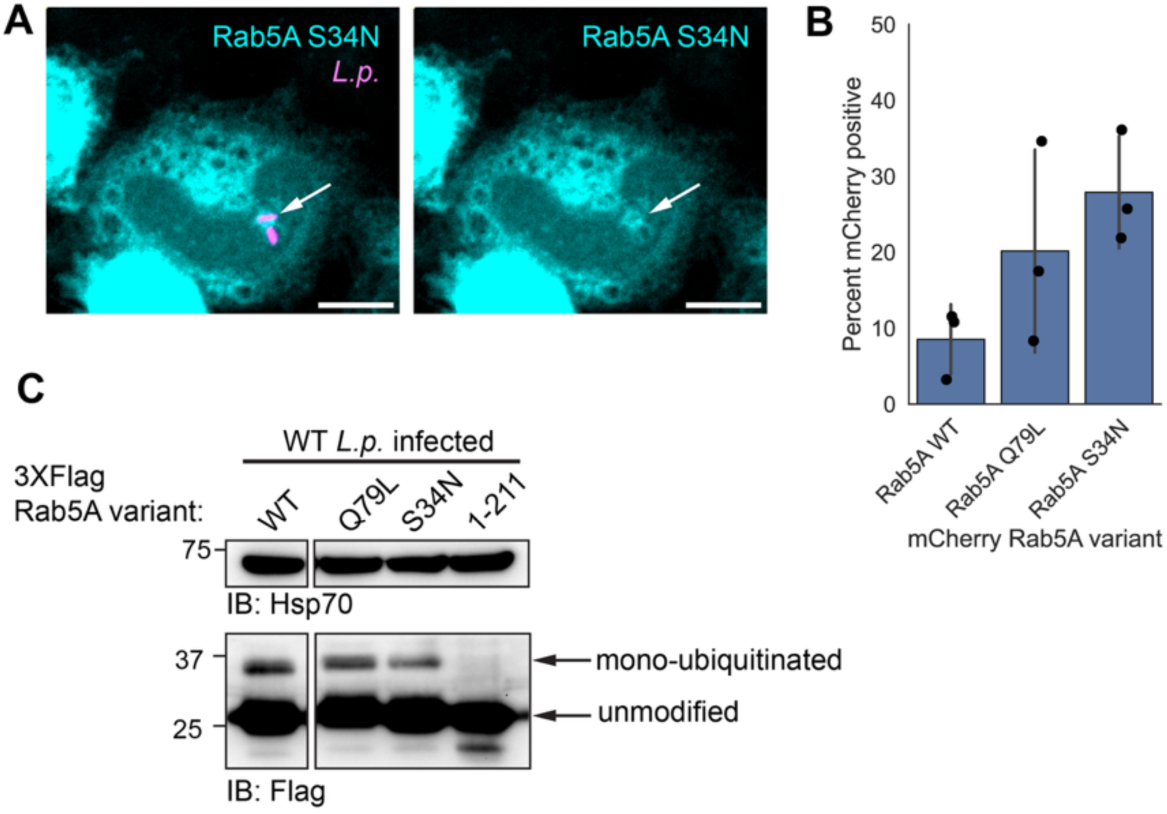
(related to Figure 1): Rab5 nucleotide binding mutants are recruited to the WT LCV and ubiquitinated during infection. (A) Representative image of mCherry Rab5A S34N recruitment to the WT LCV at 4hpi in HeLa FcγR cells. Scale bar = 10 μm. (B) Quantification of recruitment of the indicated mCherry fusion Rab5 variant to the WT LCV at 4hpi. Three biological replicates, N = 3, 25-50 LCVs analyzed per condition per replicate. (C) Western blot analysis of whole cell lysates prepared from HEK293T FcγR cells transiently transfected with the indicated Flag-tagged Rab5A variant and infected with *L.p.* WT for 5 hours.

**Figure S2.**
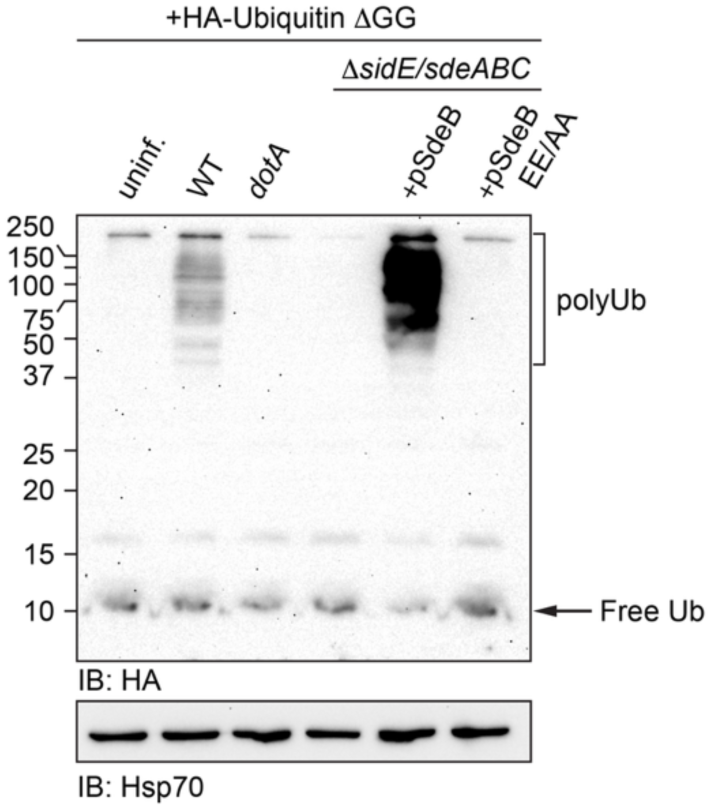
(related to Figure 2): SdeB EE/AA is unable to catalyze PR-ubiquitination. Western blot analysis of HA-Ub ι1GG conjugation in whole cell lysates prepared from HEK293T FcγR cells infected with the indicated *L.p.* strain for 1 hour.

**Figure S3:**
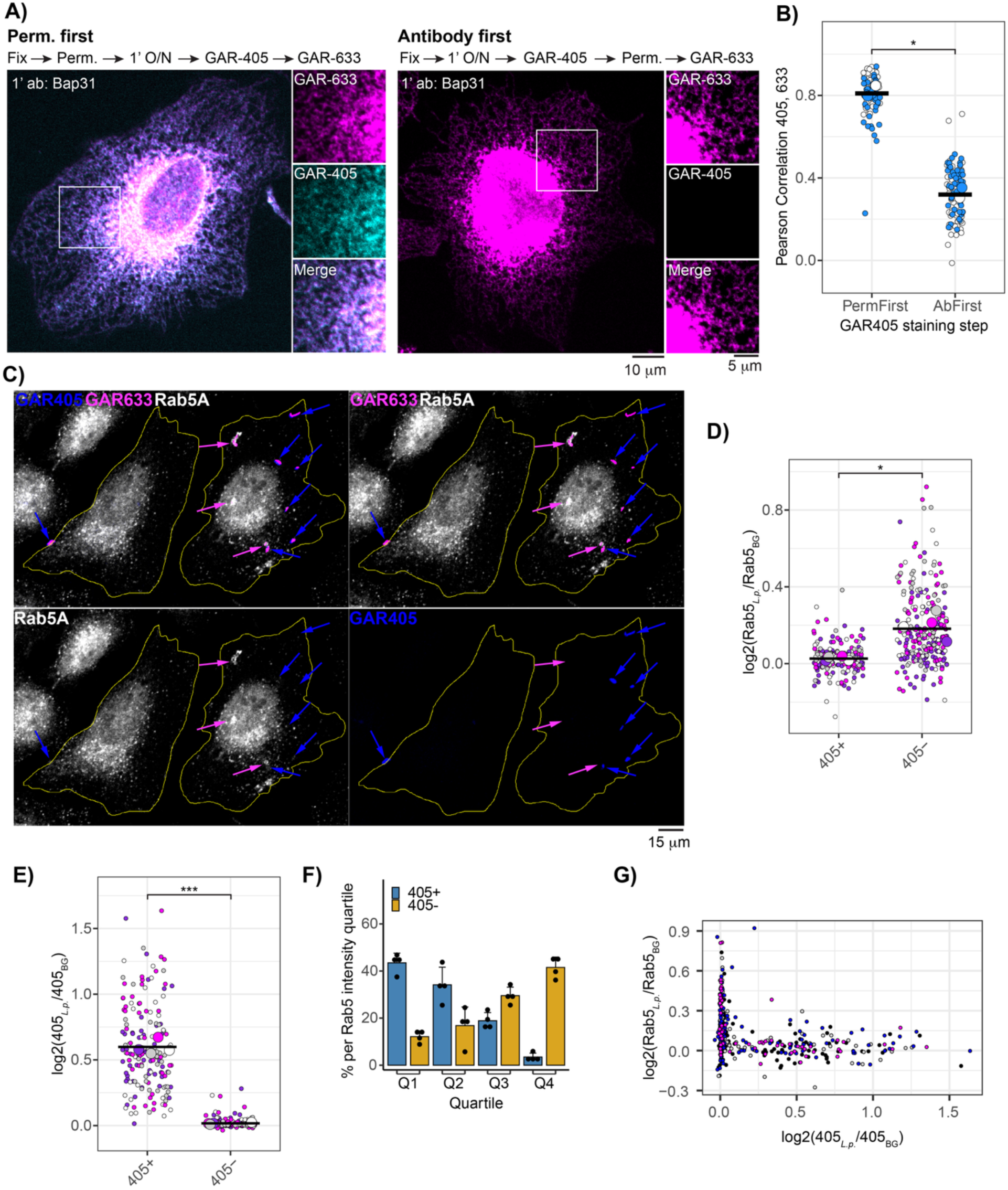
Validation of dual-stain method for detection of extracellular bacteria. (A) Representative images of cells permeabilized either before or after Bap31 primary antibody and GAR-405 staining. (B) Pearson correlation of GAR-633 and GAR-405 signal in cells prepared as in flowchart shown in A. N=2, 50 cells analyzed per replicate per condition. Welch’s two-sample t-test, p = 0.006. (C) Representative images of cells infected with *L.p.* WT and processed using dual stain method for the *L.p.* opsonization antibody, as well as immunofluorescence analysis of endogenous Rab5. Double stained (extracellular) bacteria are indicated by blue arrows, whereas single stained (intracellular) bacteria are indicated by magenta arrows. (D) Quantification of normalized Rab5 intensity at either double (405+) or single (405-) stained bacteria. N = 4, ∼30-90 bacteria scored per replicate per condition. Welch’s two sample t-test, p = 0.0127. (E) Measurement of normalized GAR405 intensity at bacterial cell bodies for same dataset described in D. Welch’s t-test, p = 0.0002. (F) Distribution of Rab5 intensity values for dataset described in D. For each replicate, data was pooled, and the percentage of 405+ versus 405- bacteria falling into each quartile was tabulated. (G) Normalized GAR405 intensity vs. normalized Rab5 intensity for dataset described in D. Note that very few datapoints show strong signal for both Rab5 and GAR405. Color represents biological replicate (N=4), 418 total bacteria scored.

**Figure S4.**
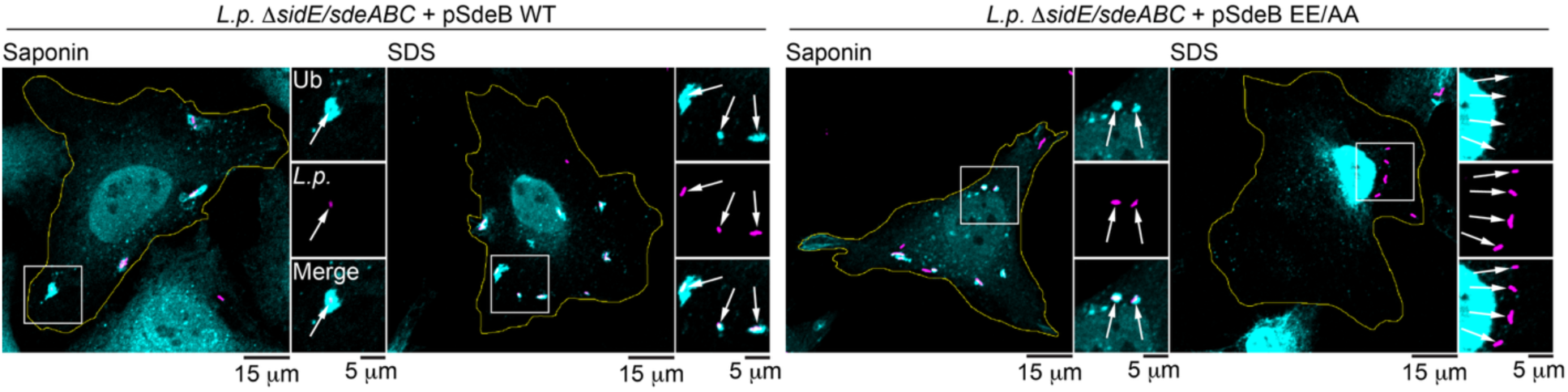
(related to Figure 5): Extended representative images of ubiquitin detergent resistance. Endogenous ubiquitin staining in cells infected with the indicated strain and permeabilized with either saponin or SDS at 1hpi.

**Figure S5.**
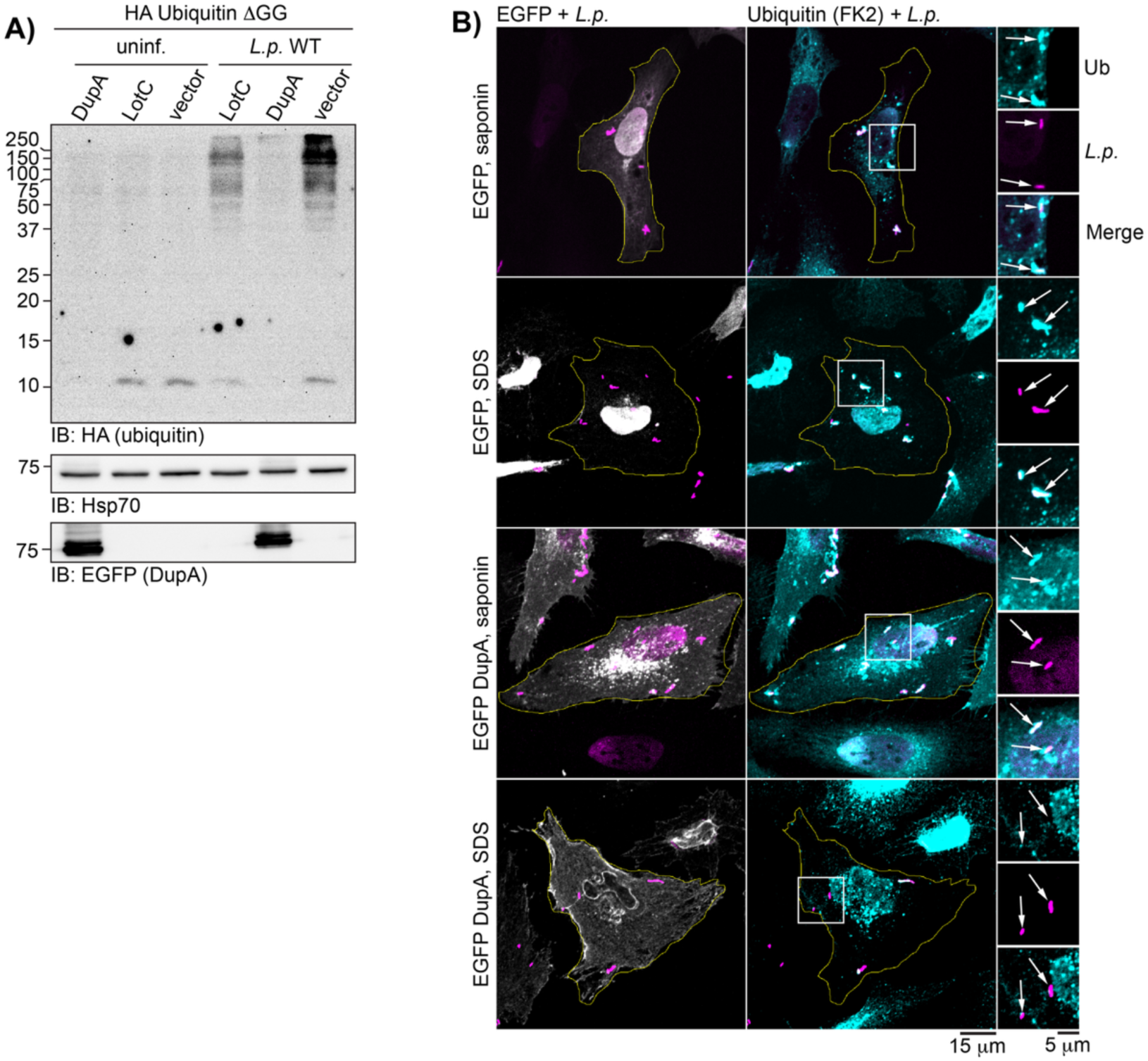
(related to Figure 6): Ectopically expressed EGFP-DupA is active against PR-ubiquitin conjugates. (A) Western blot analysis of lysates from cells transfected with HA-ubiquitin ι1GG and the indicated construct (LotC is a canonical ubiquitin ligase effector), and either left uninfected or infected with *L.p.* WT. (B) Representative images of endogenous ubiquitin staining in cells transfected with either EGFP alone or EGFP-DupA, infected with *L.p.* WT, and permeabilized with either sapoinin or SDS. For all infection experiments, cells were fixed or lysed at 1hpi.

**Figure S6.**
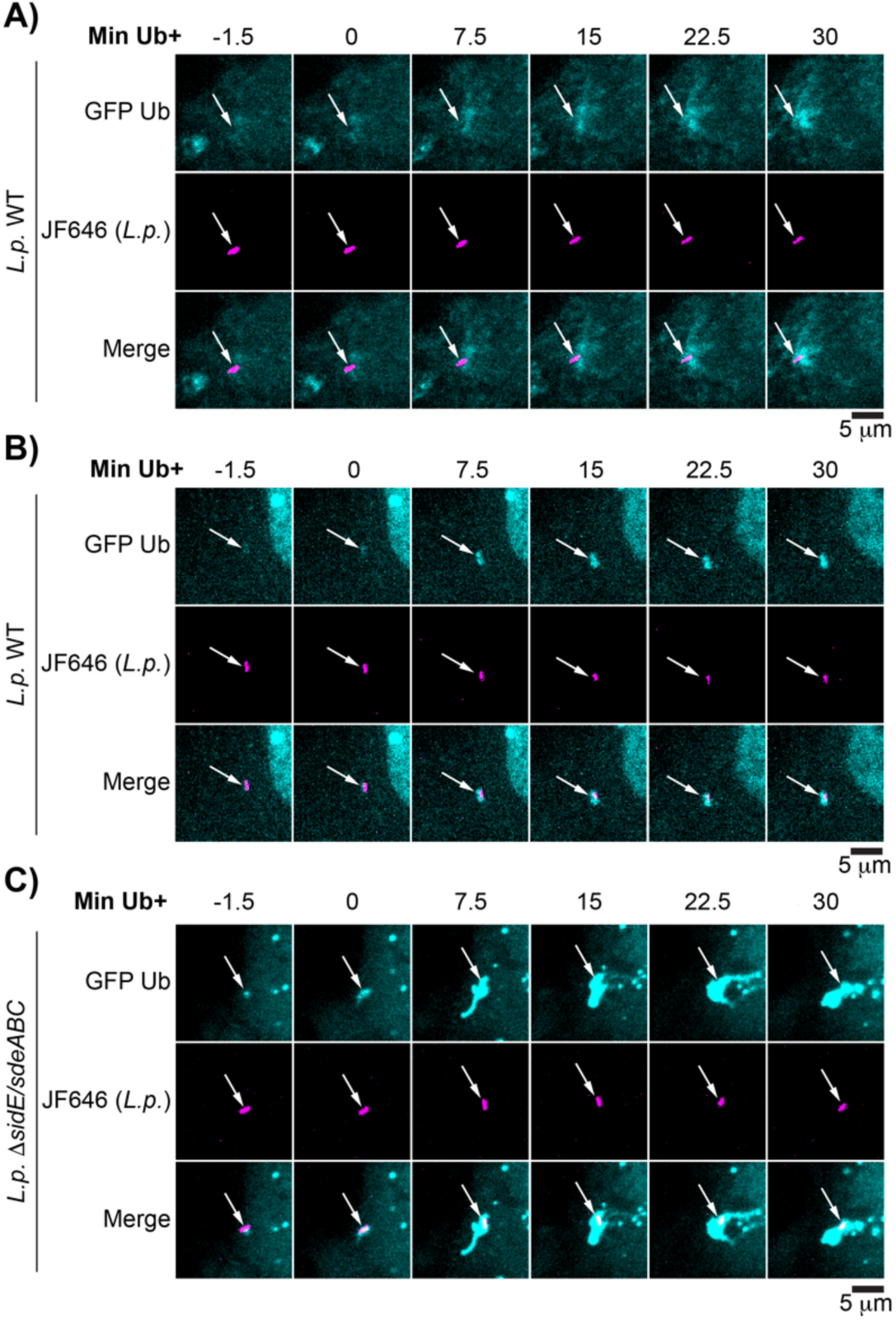
(related to Figure 7): Uncommon LCV-associated ubiquitin morphology patterns observed during live imaging. Example cases of (A) diffuse ubiquitin localization throughout imaging at WT LCV, (B) compact ubiquitin localization at WT LCV, and (C) highly dynamic compact-to-expansive ubiquitin morphology at the SidE family knockout strain LCV.

## Supplemental movies S1-4

See Methods for sample preparation and image acquisition. Scale bar length corresponds to 10 μm.

## Notes

### Competing Interest Statement

The authors have declared no competing interest.

### Summary of Updates

Additional data has been added regarding the role of the nucleotide binding state of Rab5 in recruitment to the LCV and ubiquitination during infection (see figure S1). The discussion has been expanded to include further detail about the literature regarding Legionella and the endolysosomal system. During data organization, we realized that the representative images for SDS permeabilized WT infected cells for 1- and 2- hours post-infection had been inadvertently swapped in figure 6A. While this does not affect the conclusions of the study, we sincerely apologize for the error, and have corrected it in this revised version. Finally, in recognition of her contributions to the revised version, Puspangana Singh has been added as an author.

